# Mitochondrial dysfunction rapidly modulates the abundance and thermal stability of cellular proteins

**DOI:** 10.1101/2022.10.27.514032

**Authors:** Carina Groh, Per Haberkant, Frank Stein, Sebastian Filbeck, Stefan Pfeffer, Mikhail M. Savitski, Felix Boos, Johannes M. Herrmann

**Affiliations:** Cell Biology, University of Kaiserslautern, 67663 Kaiserslautern, Germany; Proteomics Core Facility, EMBL Heidelberg, 69117 Heidelberg, Germany; ZMBH, University Heidelberg, 69120 Heidelberg, Germany; Department of Genetics, Stanford University, California, USA

**Keywords:** Mitochondria, Proteasome, Protein folding, Proteostasis, Thermal proteome profiling, Stress response

## Abstract

Cellular functionality relies on a well-balanced, but highly dynamic proteome. Dysfunction of mitochondrial protein import leads to the cytosolic accumulation of mitochondrial precursor proteins which compromise cellular proteostasis and trigger the mitoprotein-induced stress response. To dissect the effects of mitochondrial dysfunction on the cellular proteome as a whole, we developed pre-post thermal proteome profiling (ppTPP). This multiplexed time-resolved proteome-wide thermal stability profiling approach with isobaric peptide tags in combination with a pulsed SILAC labeling elucidated dynamic proteostasis changes in several dimensions: In addition to adaptations in protein abundance, we observed rapid modulations of the thermal stability of individual cellular proteins. Strikingly, different functional groups of proteins showed characteristic response patterns and reacted with group-specific kinetics, allowing the identification of the functional modules that are relevant for mitoprotein-induced stress. Thus, our new ppTPP approach uncovered a complex response network that orchestrates proteome homeostasis in eukaryotic cells by time-controlled adaptations of protein abundance and protein stability.

## Introduction

The eukaryotic proteome consists of thousands of different proteins. Changes in the developmental, environmental or metabolic conditions can induce a considerable remodeling of the proteome. Even seemingly small changes can have pronounced effects. For example, the replacement of the carbon source glucose by galactose in yeast cultures changes the expression of more than 25% of all genes at least twofold, even though galactose catabolism *per se* requires only the function of five additional enzymes (Ronen & Botstein, 2006). By complex remodeling of the proteome network, cells apparently strive for a maximum of functional performance and stability, but the molecular mechanisms underlying these transitions are only poorly understood.

Protein homeostasis (proteostasis) depends not only on the abundance of each cellular protein but also on their specific folding, membrane topology, cellular location, and interaction partners as well as on diverse posttranslational modifications. An elaborate quality control system, also termed proteostasis network, regulates the synthesis, folding, transport and degradation of proteins (Elsasser *et al*, 2022; Hipp *et al*, 2019; Sala *et al*, 2017). It relies on molecular chaperones that facilitate and stabilize protein folding (Hartl *et al*, 2011) and the ubiquitin-proteasome system (UPS) to recognize and remove damaged or surplus proteins (Dikic, 2017). In addition, a multitude of factors controls the spatial organization of the proteome, either by facilitating protein insertion into and translocation across membranes (Wickner & Schekman, 2005; Wu & Rapoport, 2021) or by concentrating proteins in membrane-less condensates (Alberti & Hyman, 2021; Sontag *et al*, 2017).

When facing a proteostasis challenge, cells activate stress response pathways that alter the composition of the proteome through changes in protein synthesis or degradation. These changes in protein abundance under different conditions can be determined at high resolution by mass spectrometry (Aebersold & Mann, 2016). However, changes in protein abundance cannot reflect the functional state of a protein, which often depends on its folding state, posttranslational modifications or binding partners. These features remain hidden in classical expression proteomics studies. This is particularly true when examining short timescales, such as the first hours after cells face a proteotoxic challenge. We therefore sought to establish a new method that provides time-resolved and proteome-wide insights into the functional state of proteins under dynamically changing stress conditions. Thermal proteome profiling (TPP) makes use of the cellular thermal shift assay (Martinez Molina *et al*, 2013) and represents the multiplexed quantification of non-denatured protein fractions as function of temperature (Franken *et al*, 2015; Mateus *et al*, 2020; Savitski *et al*, 2014). Thus, TPP provides a proxy for the thermal stability of each protein *in vivo*. This powerful strategy is perfectly suited for the detection of specific subtle changes, for example in order to identify an intracellular binding site of a chemical compound (Mateus *et al*, 2022) but also to study global changes such as the proteome-wide effects of a specific posttranslational modification (Vieitez *et al*, 2022). On the other hand, a pulsed labeling approach through the replacement of amino acids by isotopes of different mass in a growing culture (pulsed stable isotope labeling by amino acids in cell culture, pulsed SILAC) can be used to monitor proteome changes of pre-existing and newly synthesized proteins at the same time (de Godoy *et al*, 2008; Ong *et al*, 2002). This method proved to be very powerful to determine the import, assembly and degradation of mitochondrial proteins (Bogenhagen & Haley, 2020; Saladi *et al*, 2020; Schafer *et al*, 2022). In this study, we combined for the first time pulsed SILAC labeling with a two-dimensional TPP approach (2D-TPP) (Becher *et al*, 2018) which we termed pre-post-TPP (ppTPP). With this, we were able to monitor in parallel the time-sensitive variations in abundance and stability of mature and newly synthesized proteins as adaptation to a specific stress-inducing insult.

We chose the mitoprotein-induced stress response (Boos *et al*, 2020; Topf *et al*, 2019) to evaluate the potential of this method. Mitochondria consist of hundreds of proteins which are synthesized in the cytosol as precursor proteins that are post-translationally imported into the organelle using the translocase of the outer membrane (TOM complex) as common entry gate (Araiso *et al*, 2019; Chacinska *et al*, 2009). Mitochondrial dysfunction can impair protein import causing the cytosolic accumulation of mitochondrial precursor proteins. Such precursor accumulation may occur transiently during development or upon metabolic adaptations, or as a chronic state during aging and age-related diseases (Bauer & Neupert, 2001; Cenini *et al*, 2016; Coyne & Chen, 2018; Devi *et al*, 2006; Eckl *et al*, 2021; Franco-Iborra *et al*, 2018; Li *et al*, 2010; Tsuboi *et al*, 2020).

Induction of a slowly imported clogger protein from a galactose-inducible promoter competitively inhibits mitochondrial protein import and serves as an elegant model to study the consequences of cytosolic precursor proteins in cells that otherwise contain fully functional mitochondria (Fig. 1A) (Boos *et al*, 2019; Weidberg & Amon, 2018). Cells tolerate this situation well and react with a multi-level response termed unfolded protein response activated by mistargeting of proteins (UPRam) (Wrobel *et al*, 2015). This stress response is characterized by adaptive reactions, including the induction of the heat shock response, increased levels of the UPS and the repression of protein synthesis, in particular of translation products destined for mitochondria (Boos *et al*., 2019; Wrobel *et al*., 2015). But this situation also triggers mitochondria-specific reactions to remove the stalled clogger protein from the TOM complex accomplished by an interplay between a specific AAA-ATPase and proteasome-mediated proteolysis (Mårtensson *et al*, 2019; Shakya *et al*, 2021; Weidberg & Amon, 2018).

**Fig. 1.**
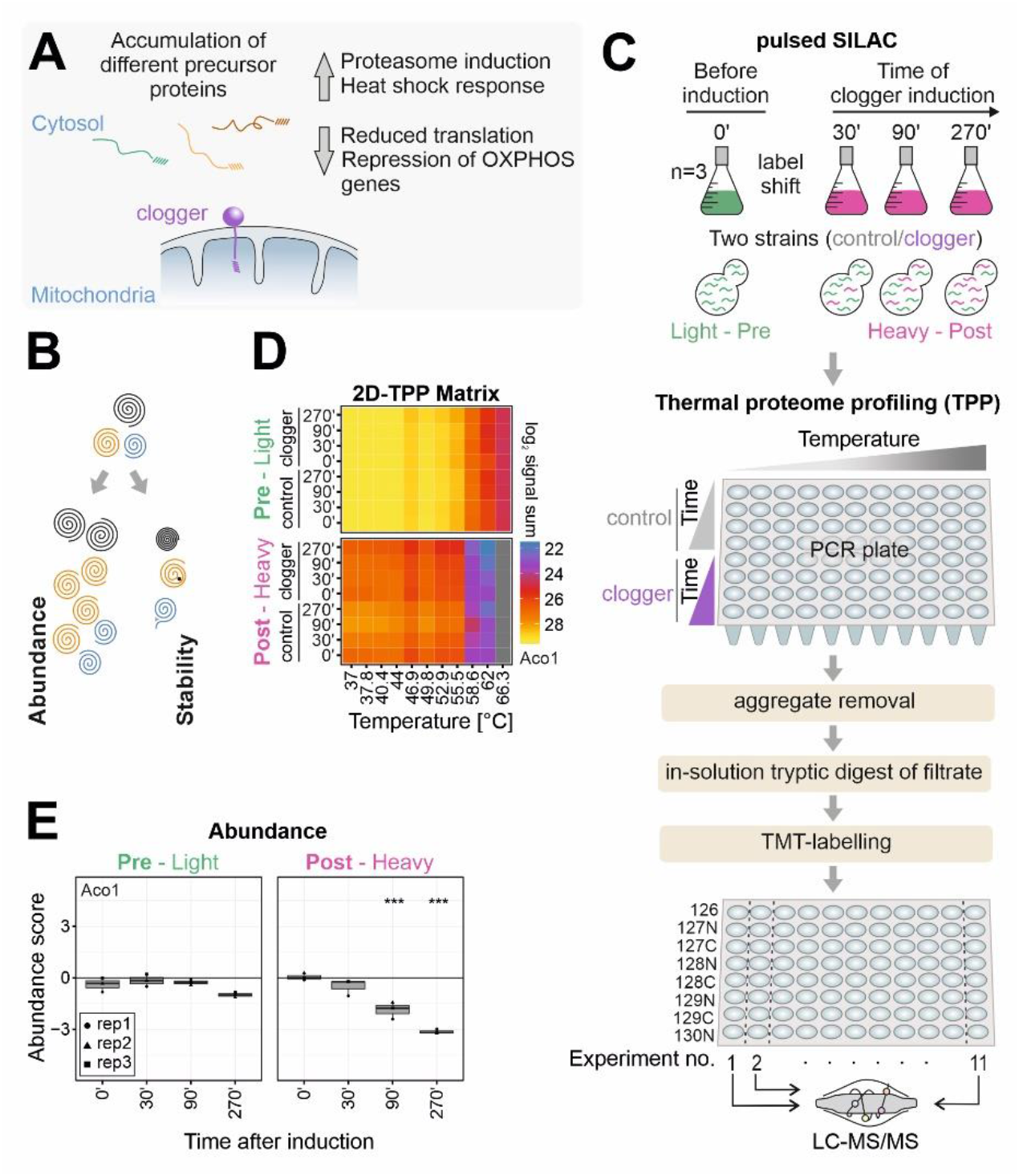
The combination of pulsed SILAC and TPP (ppTPP) allows it to measure stress-induced effects on protein abundance and thermal stability for mature and newly synthesized proteins in one multiplexed approach. **A**. Clogger induction triggers the mitoprotein-induced stress response. **B**. The proteome adapts to stress by changing the abundance and thermal stability of proteins. **C**. Workflow of the ppTPP approach. For pulsed SILAC experiment, clogger and control cells were grown in light-isotope containing respiratory medium until mid-exponential growth phase. Clogger was induced by addition of 0.5% galactose. Simultaneously, medium was switched from light to heavy isotope containing medium. Cells were harvested after different timepoints and heated to eleven different temperatures and lysed with NP40 and zymolyase. Non-soluble proteins were removed by filtration, and the remaining soluble fractions were labeled with isobaric mass tags (TMT) for protein quantification. Different conditions of each temperature were combined in single TMT-experiments, which were analyzed by LC-MS/MS. **D**. Representation of the 2D-TPP matrix. Shown is the signal sum for different times after clogger induction for proteins synthesized before (upper panel) and after (lower panel) induction in the control and clogger strain for each temperature. Mitochondrial aconitase (Aco1) is shown as representative example. See Table S1 for details. **E**. Pulsed SILAC allows to detect variations for different protein states. Shown is abundance score (clogger/control) for different times after clogger induction for proteins synthesized before (left panel) or after (right panel) induction. Mitochondrial aconitase (Aco1) is shown as representative example. Asterisks indicating significant pvalue (***, <0.001).

Mitoprotein-induced stress response seems perfectly suited to explore the potential of ppTPP for several reasons: (i) despite of the immediate and stark response of the cell to clogger induction, cells (and also their mitochondria) stay functional for several hours and maintain their viability; (ii) effects on lipids and metabolites are only secondary through changes in the proteome and, hence, are expected to be much less pronounced than upon heat shock or oxidative stress conditions which directly affect these molecules; (iii) the defined site of action (the clogged mitochondrial TOM complex) allows to distinguish immediate effects of the stress on mitochondrial proteins as well as secondary effects in the cytosol and other ‘distant’ cellular compartments.

The multidimensional analysis of the ppTPP approach revealed that the cellular proteome reacts on different levels, including various time-dependent changes in protein abundance and stability. Strikingly, different groups of proteins show highly distinct response patterns and react with subgroup-specific kinetics. Many of these patterns would not have been detectable or discernible by analyzing only abundance, demonstrating the power of ppTPP to discover novel pathways, especially those that act on short timescales.

## Results

### The combination of pulsed SILAC and 2D-TPP enables the proteome-wide profiling of thermal stability and abundance of mature and newly synthesized proteins upon stress induction

The impact of individual aggregation-prone model proteins on cellular fitness and neurodegeneration is well documented. The mitoprotein-induced stress response is arguably more complex as it is elicited by the simultaneous accumulation of hundreds of mitochondrial precursor proteins in the cytosol which exert a complex pattern of effects in the cell (Fig. 1A)(Boos *et al*., 2019; Schafer *et al*., 2022; Shakya *et al*., 2021; Wrobel *et al*., 2015; Xiao *et al*, 2021). The galactose-controlled induction of the slowly imported model protein cytochrome *b_2_*(1-167)-dihydrofolate reductase (*b*_2_-DHFR, here also referred to as ‘clogger’) in cells of the yeast *Saccharomyces cerevisiae* proved to serve as a powerful model system for which detailed transcriptome and proteome data are available, demonstrating a complex cellular response with mitochondrion-specific but also generic elements (Boos *et al*., 2019; Weidberg & Amon, 2018). As a control for the clogger protein we employed galactose-inducible cytosolic DHFR to address galactose induced changes that are independent of mitochondrial protein import (Fig. S1A). Expression of the clogger, but not of the DHFR control, slows down cell growth (Fig. S1B, C).

To elucidate the clogger-induced adaptations on the abundance and thermal stability (Fig. 1B, S1D) of proteins synthesized before or after onset of clogger expression, we combined 2D-thermal proteome profiling (Becher *et al*., 2018) with pulsed SILAC in an integrated approach which we termed pre-post-TPP (ppTPP, Fig. 1C, S1E). Cells with plasmids for induction of the clogger or the DHFR control were grown to mid-exponential phase on lactate medium containing ^14^N_4_-^12^C_6_-arginine and ^14^N_2_-^12^C_6_-lysine (light, pre induction). Then, cells were isolated and transferred to lactate plus galactose medium containing ^15^N_4_-^13^C_6_-arginine and ^15^N_2_-^13^C_6_-lysine (heavy, post induction). After 30, 90 and 270 min of expression of clogger or control, aliquots were taken, and cells were subjected to a 3 min incubation at 11 different temperatures ranging from 37°C to 66.3°C. After lysis with the mild detergent NP40 and the cell wall-digesting enzyme zymolyase, non-soluble proteins were removed, and NP40-soluble proteins were identified by multiplexed quantitative mass spectrometry based on isobaric tandem mass tag (TMT) labelling. To this end, the individual samples of each time point were TMT-labelled, multiplexed and measured in a single mass spectrometry-based experiment to determine changes in thermal stability and abundance along the time of clogger-induction (Fig. 1D).

Using the DHFR strain as control to calculate fold changes (FC), the matrix was condensed to two measures: (i) the abundance (corresponding to the average changes in the two lowest temperatures) and (ii) stability score (corresponding to changes remaining at higher temperatures after correcting for abundance changes) were calculated using published procedures (Fig. S2A)(Franken *et al*., 2015; Mateus *et al*., 2020).

The protein abundances resulted in generally stable levels of light-encoded proteins. In contrast, the newly synthesized heavy-encoded proteins showed much stronger protein-specific responses to clogger expression, robustly demonstrating that the SILAC labelling reliably distinguished pre-existing proteins from those synthesized after clogger expression. For example, newly synthesized aconitase (Aco1), like many mitochondrial proteins, was steadily diminished upon clogger induction compared to control, while the levels of pre-existing Aco1 were barely influenced (Fig. 1E).

Thus, the ppTPP procedure allows it to systematically assess the proteome-wide abundance and thermal stability of nascent and mature proteins before and after onset of stress conditions in one single experiment.

### Stress influences the abundance and the thermal stability of different groups of proteins in characteristic patterns

Do the changes in abundance correlate with those in thermal stability (to which we refer as stability in the following for simplicity)? To answer this question, we calculated pairwise Pearson correlation coefficients for all different samples (Fig. 2A) and plotted the changes in stability against those in abundance (Fig. S2B). This showed that changes in the stability correlated well with stability changes, and changes in abundance with abundance changes throughout the different samples and timepoints. The correlations increased over time of clogger expression both for abundance and for stability, indicating that increasing stress conditions shape a more and more defined and consistent pattern of proteome reorganization (Fig. 2A, arrows). However, changes in the abundance and in the stability of proteins represent distinct, non-correlating dimensions. The correlated variables are fundamentally different and show almost no relationship, or only a slight negative correlation for preexisting, but not for newly synthesized proteins.

**Fig. 2.**
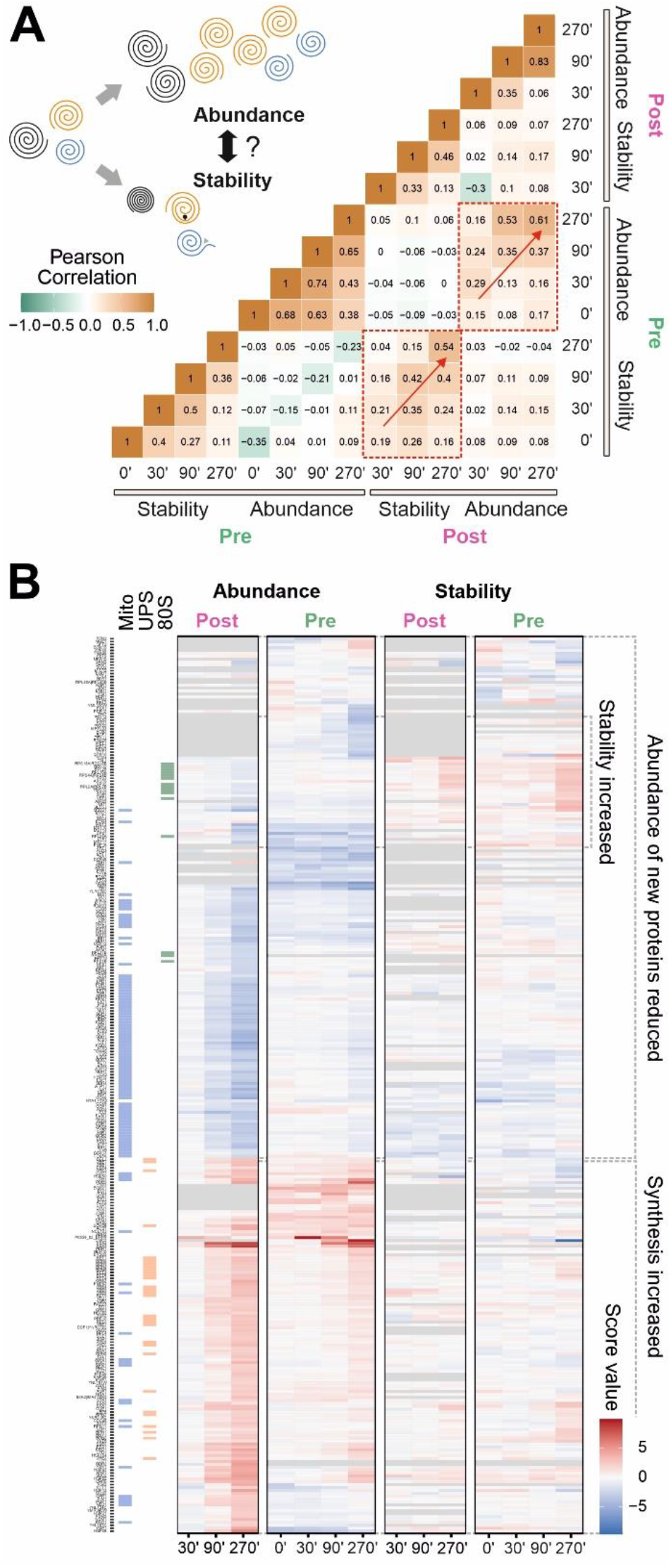
Stress-induced changes in abundance and stability do not correlate. **A**. Pearson correlation coefficients of all pairwise proteins were calculated using the data generated during ppTPP (clogger vs. control). Whereas the different abundance and stability scores correlate with each other, abundance does not or very poorly correlate with stability (see also Fig. S2). Red arrows are indicating an increasing correlation with prolonged stress. **B**. Clustering of stability and abundance of significantly changing proteins. Score values of the different samples (clogger vs. control) are shown and the distribution of mitochondrial, proteasomal and ribosomal proteins is indicated on the left. See also Table S1 for full dataset.

A heatmap showing the score values of proteins which responded significantly to clogger induction showed the most prevalent changes were those of peptide abundance of newly synthesized proteins (Fig. 2B). Many mitochondrial proteins were reduced upon clogger expression (Fig. 2B, see proteins marked in blue on left) as a result of their competition with the clogger for mitochondrial import sites. Conversely, proteins of the UPS (Fig. 2B, marked in orange on left) showed the opposite trend and were increased upon clogger expression, consistent with the characteristic proteasome induction previously reported on a gene expression level (Boos *et al*., 2019). Strikingly, also the stability changes showed consistent effects on specific groups of proteins. For example, the stability of many cytosolic ribosomal proteins increased upon clogger expression (Fig. 2B, marked in green on left). Stability, in contrast to abundance, affected both newly synthesized proteins and proteins that were present before clogger was expressed. Notably, with our SILAC-combined TPP, it is possible to analyze the direct structural and hence functional effect of the clogger on freshly synthesized mitochondrial proteins, which would be hidden when using TPP only (Fig. S2C).

Our data demonstrate that abundance and stability changes occur largely independently from each other and are remodeled during mitoprotein-induced stress in a pattern that characteristically and consistently reflects the physiological role of different protein groups.

### Mitoprotein-induced stress strongly diminishes the synthesis of mitochondrial proteins

First, we studied the effect of clogger expression on the abundance of proteins. The signals of light peptides diminished only slightly in the course of 4.5 hours whereas the heavy peptides increased strongly (Fig. 3A). Compared to control cells, clogger expression globally reduced protein synthesis, in particular upon prolonged clogger induction (Fig. 3B, S3A). Thus, the rates of protein synthesis (Fig. S3B) were significantly reduced upon mitoprotein-induced stress (Fig. 3C, D).

**Fig. 3.**
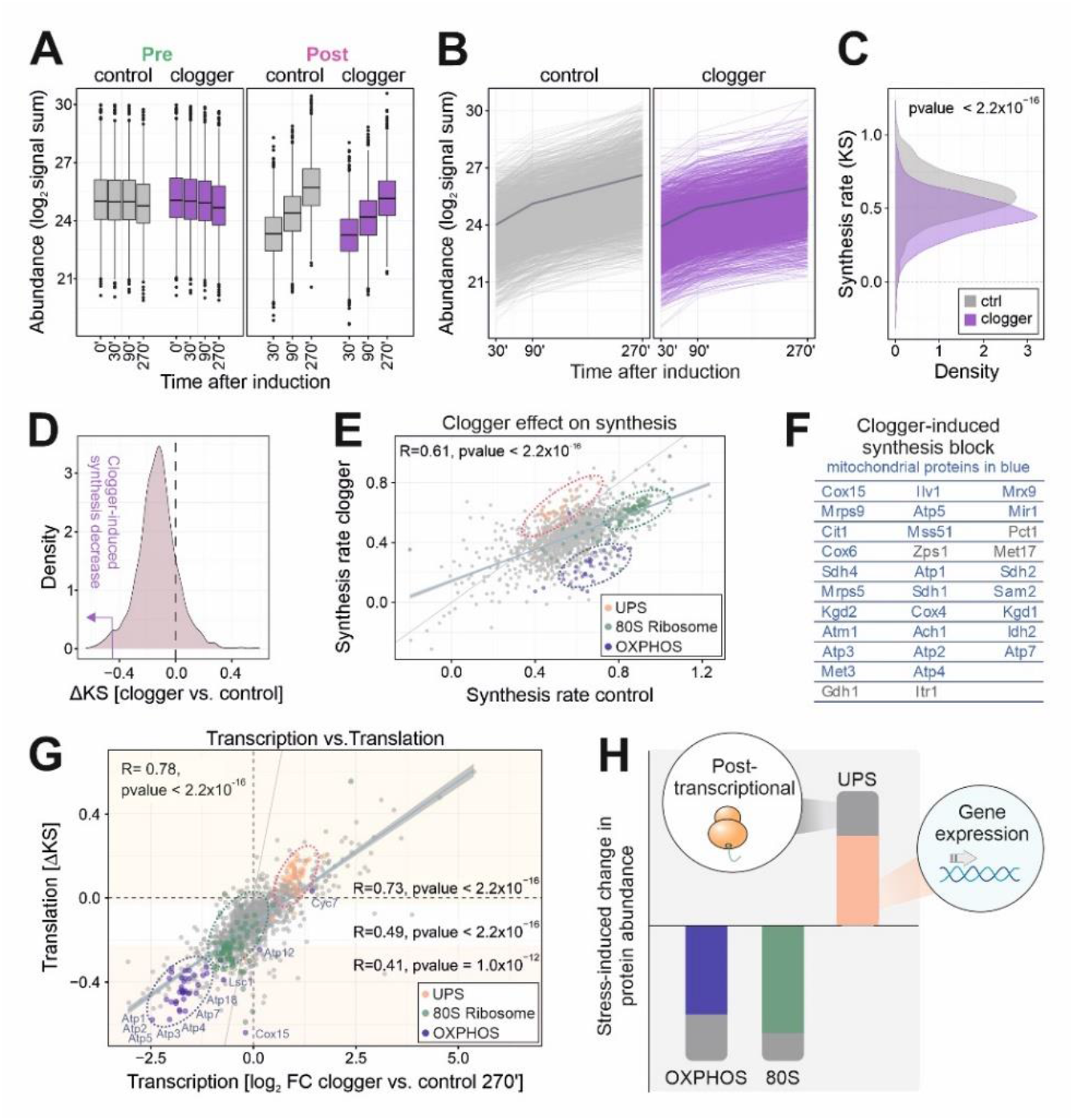
The stress-response affects protein synthesis by class-specific transcriptional and post-transcriptional adaptations. **A**. Total protein signal sums show the increase of newly synthesized proteins over time (right panel) in contrast to preexisting proteins shown in the left panel. y-axis shows signal intensities of the protein abundance. Line represents the median, box represents the interquartile range, and whiskers of the box represent the 5^th^ and 95^th^ percentiles. **B**. Signal sums of newly synthesized (heavy) peptides in the absence and presence of clogger. The dark line indicates the mean value over all proteins. **C**. Density plot showing the synthesis rate (KS) along the 270 min course of clogger induction of stressed and control cells. Synthesis rate was calculated on the basis of heavy peptide intensities. Two-sample Kolmogorov-Smirnov test demonstrates a significant difference between stressed and unstressed cells, revealing lower synthesis rates upon clogger expression. **D**. Clogger expression reduces synthesis of most proteins. Density plot of ΔKS, calculated by the ratio of the synthesis rate (KS) of the clogger and control cells. Purple arrow indicating the area used for outlier analysis. **E**. The synthesis rates of proteins in the presence or absence of clogger were correlated. Proteins of the UPS, the 80S ribosome and the OXPHOS were labeled. R, Pearson correlation coefficient **F**. List of proteins with most pronounced clogger-induced inhibition of synthesis rates. Mitochondrial proteins are indicated in blue. **G**. The clogger-induced changes in transcription after 270 min (Boos *et al*., 2019) were compared to that in protein synthesis rates (clogger vs. control) showing an overall correlation of 0.78. Pearson correlation coefficient was additionally calculated between transcription and different translation rates: the upper 20%, lower 20% and middle 60% of the dataset. The names of some proteins with strong clogger-induced synthesis blocks were indicated. Proteins of the UPS, the 80S ribosome and the OXPHOS were labeled. R, Pearson correlation coefficient **H**. Mitoprotein-induced stress results in characteristic changes in protein abundance which are to a large part the result of changes in gene expression (shown in color).

When the synthesis rates with and without clogger were compared, different functional protein groups showed characteristic footprints (Fig. 3E, S3C): The clogger strongly prevented the accumulation of newly synthesized proteins of the mitochondrial OXPHOS system, either by reducing their synthesis or by inducing their immediate degradation (Fig. 3E dark blue, Fig. S3C, D). On the contrary, the synthesis of proteins of the UPS was strongly increased upon clogger expression (Fig. 3E orange, Fig. S3C, E).

Subsequently, we tested which individual proteins were most drastically diminished upon clogger expression. (Fig. 3D). Almost all of the most clogger-reduced proteins were proteins located to mitochondria (Fig. 3F). Interestingly, this group contained inner membrane proteins (Atm1, Cox15) that were found to be co-translationally imported into yeast mitochondria (Williams *et al*, 2014).

The clogger-induced changes in protein synthesis rates correlate generally well with the changes in mRNA levels (Boos *et al*., 2019). The correlation was more pronounced for abundant proteins than for lower expressed proteins (Fig. S4A-C). However, two groups of proteins deviate from this close correlation: OXPHOS components and other mitochondrial proteins are much more diminished by clogger expression than expected from their mRNA levels, presumably as a consequence of their direct competition with the clogger (Fig. 3G). On the other hand, subunits of the proteasome accumulated disproportionately more than expected from their transcript levels. This is in line with recent observations that the unfolded protein response activated by mistargeting of proteins (UPRam) directly promotes proteasome assembly by hyperactivating the biogenesis factors Irc25 and Poc4 (Kusmierczyk *et al*, 2008; Le Tallec *et al*, 2007; Wrobel *et al*., 2015). Ribosomal proteins do not show this distinct separation from the general effect (Fig. 3E, G, green). In summary, cells complement the stress-induced changes in gene expression by consistent and protein-specific regulation patterns that operate on a post-transcriptional level (Fig. 3H).

### Protein turnover supports the stress-induced remodeling of the proteome

The decline of signal intensities for light peptides over time allowed us to monitor the degradation of proteins that had been synthesized prior to clogger expression (Fig. 4A, Fig. S5A). Inhibition of mitochondrial protein import stimulated the protein degradation rate in general to some degree (Fig. 4B, C, D), potentially as a result of the increased proteasome levels. The effect of the clogger on the mitochondrial proteins (Fig. 4E) goes in line with the general trend, indicating that intra-mitochondrial protein turnover is accelerated to a specific extent if protein import is blocked. Interestingly, clogger expression increased the selective degradation of specific proteins, including some mitochondrial proteins such as the outer membrane fusion protein mitofuzin (Fzo1) and the MICOS protein Mic12 (Fig. 4D, F).

**Fig. 4.**
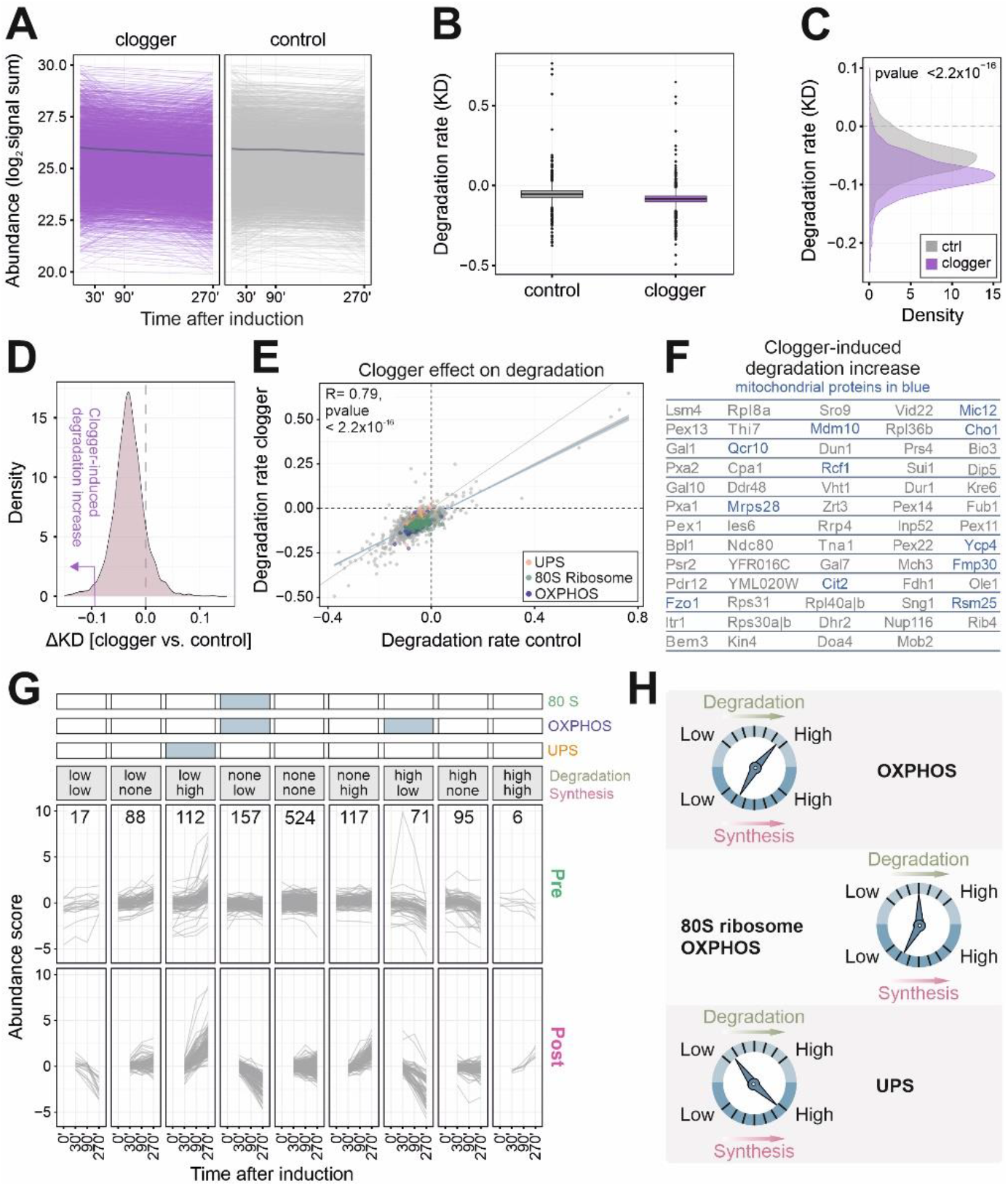
Competitive inhibition of mitochondrial protein import promotes cellular protein degradation on a large scale. **A**. Signal sums of mature (light) peptides in the presence and absence of clogger. The dark line indicates the mean value over all proteins. **B**. Box plot demonstrating the degradation rate (KD) along the 270 min course of clogger induction of stressed and control cells. Degradation rate was calculated on the basis of light peptide intensities, revealing higher degradation rates upon clogger expression. **C**. Degradation rate showing a significantly higher protein degradation of pre-existing proteins upon clogger expression. Two-sample Kolmogorov-Smirnov test was used. **D**. Density plot of ΔKD, calculated by the ratio of the degradation rate (KD) of the clogger and control cells. Purple arrow indicating the area used for outlier analysis. **E**. The degradation rates of proteins in the presence or absence of clogger were correlated, revealing only minor group-specific effects. Proteins of the UPS, the 80S ribosome and the OXPHOS were labeled. R, Pearson correlation coefficient. **F**. Proteins whose degradation was substantially increased upon clogger expression. Mitochondrial proteins are shown in blue. **G**. Classification for the different categories of abundance regulation. ΔK value of the top or bottom 20% were defined as a considerable change. Numbers are indicating proteins in respective class. Upper part demonstrating enriched protein groups in the respective classes. GO-analysis was performed using the GOrilla tool (http://cbl-gorilla.cs.technion.ac.il/), proteins of respective class were used as target set with all quantified proteins as background. Enrichment is shown with a false discovery rate [FDR] < 5% and linked to the different classes. See Table S2 for Details. **H**. Degradation and synthesis often show opposite trends, however, the changes in synthesis are much more pronounced than those in degradation.

Unexpectedly, clogger expression induced the degradation of a number of peroxisomal proteins (Pex1, Pex14, Pex22, Pxa1, Pxa2), indicating a tight crosstalk in the biogenesis of mitochondria and peroxisomes. Surprisingly, peroxisomal proteins, especially matrix proteins, were enriched in the class in which synthesis but also degradation was increased, which indicates a strong protein turnover (Fig 4G, high-high).

In many cases, though not always, protein synthesis and degradation show opposed trends (Fig. 4G): Clogger expression reduces the synthesis but stimulates the degradation of OXPHOS, to ensure a fast and effective protein clearance (Fig. S5B). On the contrary, mitoprotein-induced stress increases the production of proteasomes and at the same time the degradation of UPS components is reduced (Fig. S5C). However, ribosomal protein levels as well as specific OXPHOS proteins are mostly regulated via reduced synthesis (Fig. 4H). In summary, the selective degradation of proteins supports the clogger-induced remodeling of the proteome, however, the contribution of proteolysis is much smaller than that of altered protein synthesis.

### Stability changes reveal clogger-induced repression in protein synthesis

Since changes in abundance do not reflect structural changes of the proteome, we investigated structural proteome variations as a result of a perturbed proteostasis. The heatmap shown in Fig. 2B already revealed that clogger induction leads to a characteristic increase in the thermal stability of proteins of the cytosolic 80S ribosome (Fig. 5A). This tendency was obvious for newly synthesized ribosomal proteins as well as for proteins that had been synthesized before clogger expression (Fig. 5B-D, S6A-C). Thus, inhibition of mitochondrial protein import apparently changes the physical state of the cytosolic translation machinery, influencing thereby both preexisting (i.e. assembled) and newly synthesized ribosomal proteins.

**Fig. 5.**
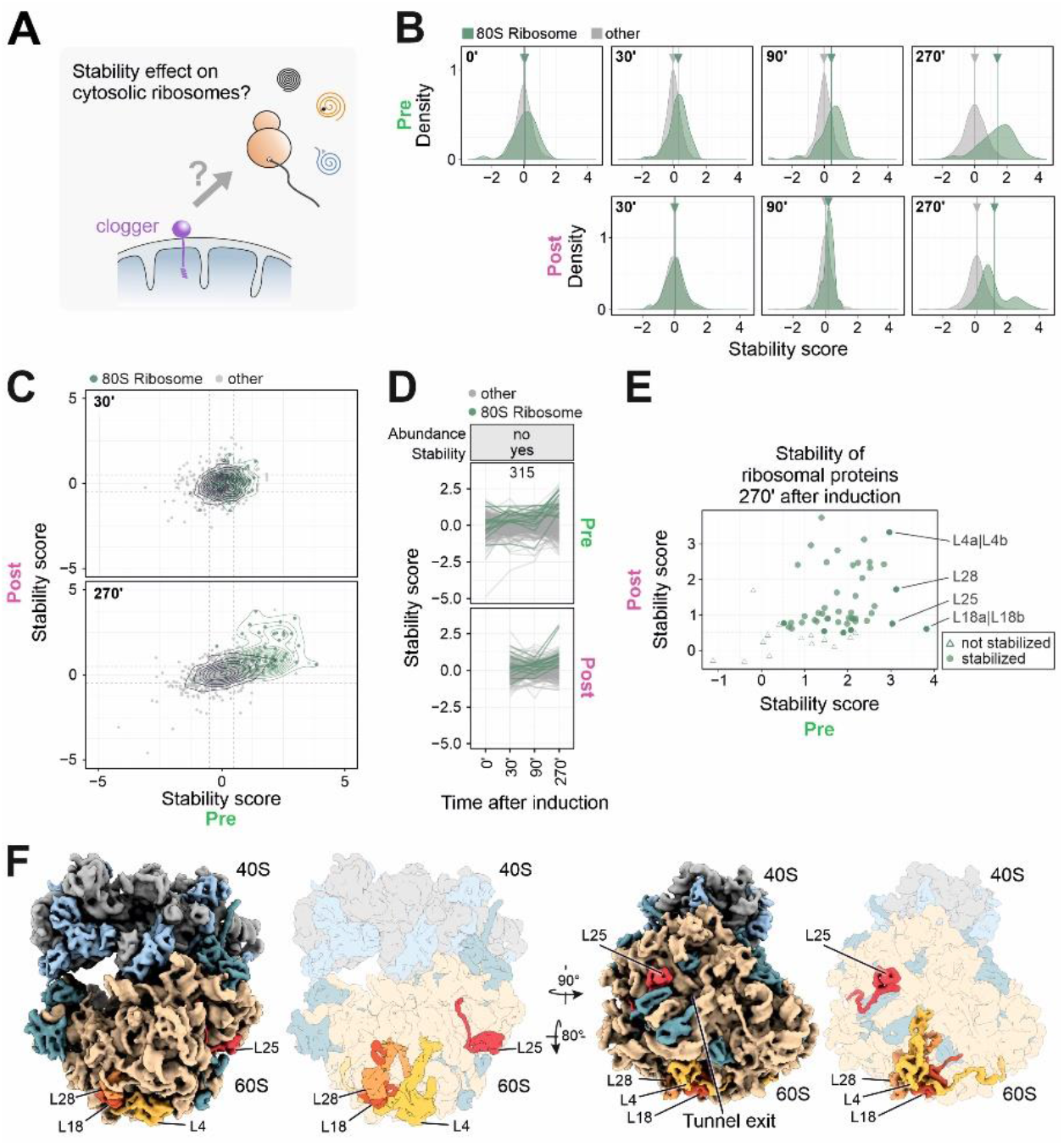
Clogger expression increases the stability of many ribosomal proteins. **A**. Potential impact of clogger-induced import inhibition on the stability of proteins of cytosolic ribosomes. **B**. The stability scores of 80S constituents that were synthesized before (pre) or after (post) clogger induction were analyzed. **C**. Stability changes (clogger vs. control) of preexisting (x-axis) and newly synthesized proteins (y-axis) were correlated. Green circles show the isobaric distribution of 80S ribosomal proteins, which show an increased stability for both protein pools over time. Black circles show the distribution of the whole proteome. **D**. Stability scores were plotted of proteins which did not change in abundance but in stability (ΔK value of the top or bottom 20% was defined as a change in abundance; a stability score below −0.5 or above 0.5 was considered as change; the numbers indicate proteins in that category). Traces of cytosolic ribosomal proteins are shown in dark green. See also Fig. S6C and Table S3 for details. **E**. Stability change of cytosolic ribosomal proteins for both preexisting and newly synthesized proteins. Four proteins of the large subunit were indicated which showed the strongest clogger-induced stabilization. (**F)**The positions of ribosomal proteins with considerably increased stability scores are shown. Simulated 3D density of an *S. cerevisiae* 80S ribosome (based on PDB-6Q8Y)(Tesina *et al*., 2019): Stabilized ribosomal proteins colored in light (SSU) and darker blue (LSU). Highly stabilized mature ribosomal proteins colored individually. Clogger expression increases the thermal stability of many proteins that are in proximity to the tunnel exit of the large ribosomal. See also Fig. S7.

Clogger-induced stabilization of mature cytosolic ribosomal proteins was most pronounced for constituents of the large subunit such as L4, L18, L25 and L28 (Fig. 5E, Fig. S6D), which are located in close proximity to the peptide tunnel exit (L25) and contribute to the structural core of the tunnel vestibule (L4, L18, L25) (Fig. 5F, Fig. S7). Also many other ribosomal proteins were stabilized, though to a lesser degree (Fig. S7). Thus, clogger induction leaves a profound footprint on the structure (and thus the function) of the ribosome which could be cause or consequence of the reduced translation rates observed when mitochondrial protein import is inhibited.

### Clogger expression alters the thermal stability of a specific set of mitochondrial proteins

In respect to their stability, mitochondrial proteins were the group of proteins that was most drastically affected by clogger expression. Clogger expression strongly destabilized mitochondrial proteins, in particular those that were synthesized after clogger induction (Fig. 6A). Since these proteins directly compete with the clogger for mitochondrial import sites, this destabilization might arise from their accumulation in the cytosol or their mislocalization to other cellular compartments (Hansen *et al*, 2018; Shakya *et al*, 2020; Weidberg & Amon, 2018; Wrobel *et al*., 2015; Xiao *et al*., 2021).

**Fig. 6.**
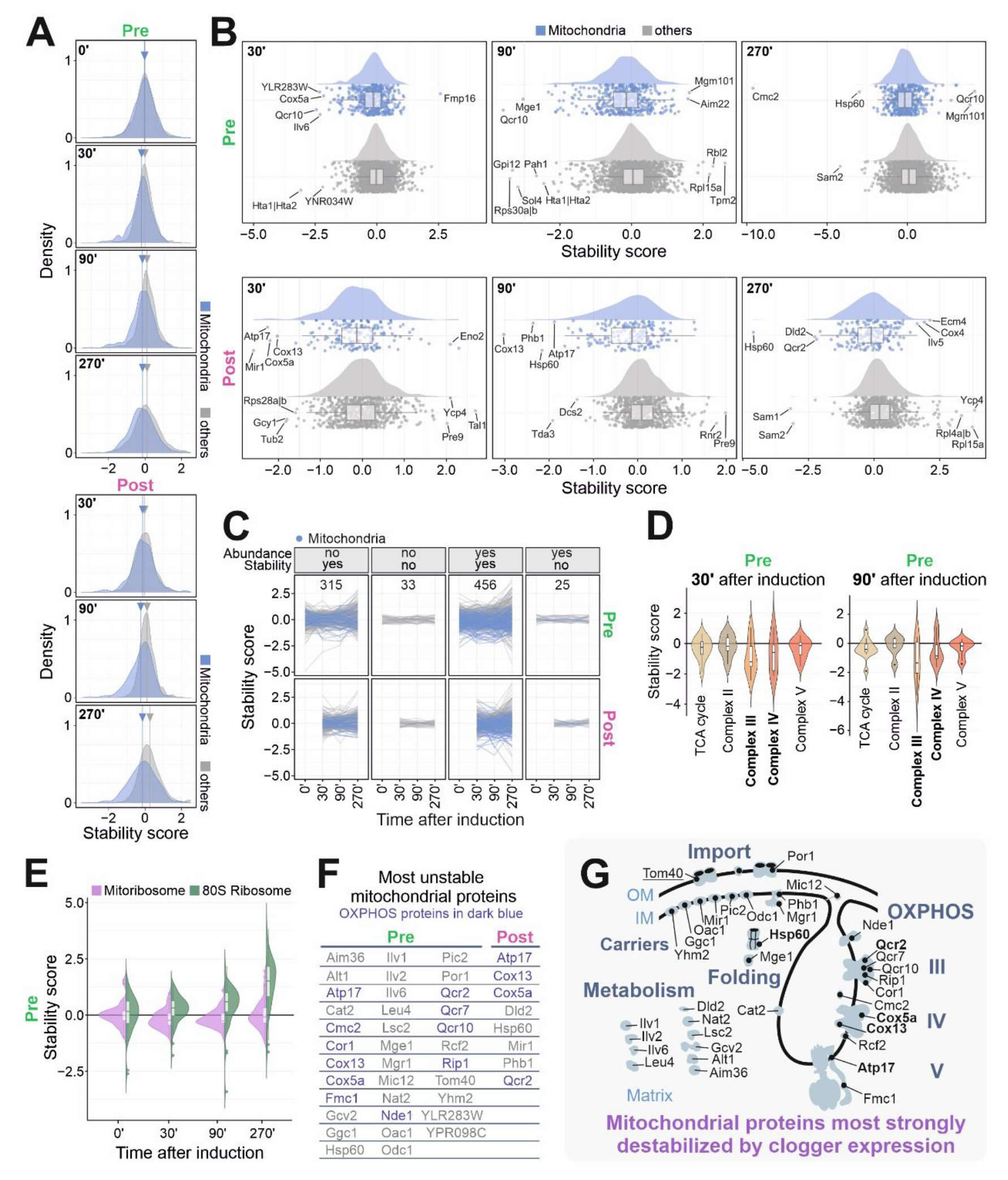
Inhibition of protein import reduces the thermal stability of many mitochondrial proteins. **A**. Kernel Density Estimation of stability scores for mitochondrial and non-mitochondrial proteins. Arrow with line indicating the mean values of the stability score, demonstrating the pronounced decrease in stability for mitochondrial proteins upon clogger expression. **B**. The clogger-induced change in thermal stability was plotted for non-mitochondrial (grey) and mitochondrial (blue) proteins. **C**. Classification according to Fig. 5D, S6C. Mitochondrial proteins are shown in blue. See Table S4 for details. **D, E**. Range of stability changes shown for different groups of proteins. See also Fig. S8. **F, G**. Shown are mitochondrial proteins which showed most severe reduction in stability scores upon clogger expression. Unique outlier proteins for every timepoint in each protein pool based on IQR criterion. OXPHOS proteins are indicated in dark purple. Proteins for which both light and heavy peptides showed severe stability changes are labelled in bold.

In addition to this rather general destabilizing effect, clogger expression caused a particularly strong effect on the stability of a small set of mitochondrial proteins, such as Atp17, Cox5a, Cox13, Hsp60 or Qcr10 (Fig. 6B, C). Interestingly, here even those proteins were destabilized that had been already present before clogger induction, suggesting that the stability of these proteins is directly or indirectly influenced by the import rate of proteins into mitochondria. Stability changes were very pronounced for the enzymes of the OXPHOS system, in particular for mature complex III and IV (Fig. 6D, S8). In contrast to the effects found for the 80S ribosome (see above), the stability of mitochondrial ribosomal proteins was not considerably affected by clogger expression (Fig. 6E). Thus, impaired protein import into mitochondria leads to considerable changes in the thermal stability of many mitochondrial proteins, including subunits of specific multisubunit complexes of the matrix and the inner membrane (Fig. 6F). Since these changes affect both newly made and pre-existing proteins, it is likely that they indicate a profound remodeling of mitochondrial energy and amino acid metabolism (Fig. 6G).

However, not only mitochondrial proteins showed an altered stability upon clogger expression (Fig. S9): For example, we noticed profound changes in the stability of the histones H2A, H3 and H4 that form the nucleosome core complex for DNA packaging (Hta1, Hta2, Hht1 and Hhf1) as well as many of the coat proteins of clathrin, COP I and COP II vesicles (Fig. S10A). Apparently, clogger-induced stress leads to a complex and comprehensive rewiring of cellular physiology, including processes such as chromatin packaging and intracellular vesicle transport.

### Stability changes as predicter of chaperone occupancy

Are changes in thermal stability suited to predict molecular consequences of clogger expression on a molecular level? To address this question, we had a closer look at the clogger-induced stability changes of molecular chaperones. Clogger expression changed the stability of specific chaperones, in particular of the cytosolic Hsp70 proteins Ssa1 and Ssa2, as well as of the mitochondrial Hsp60 chaperonin (Fig. 7A, B, Fig. S10B). The direct interaction of both chaperone groups with mitochondrial precursor proteins is well established (Cheng *et al*, 1989; Endo *et al*, 1996; Hoseini *et al*, 2016; Langer *et al*, 1992; Ostermann *et al*, 1989; Smith & Yaffe, 1991). Moreover, clogger expression strongly changed the stability of the cytosolic disaggregase Hsp104 (Fig. S10C); the latter has recently been implicated to be of particular relevance in the context of mitoprotein-induced stress (ka chaNowicka *et al*, 2021; Krämer *et al*, 2022). Thus, chaperones do not follow a clear coherent tendency in stability but rather react to clogger expression with protein-specific changes in abundance and stability. This is in contrast to what we had observed for proteins of the proteasome, the 80S ribosome or the OXPHOS system, which mainly respond in concerted group-specific patterns (Fig. 7C).

**Fig. 7.**
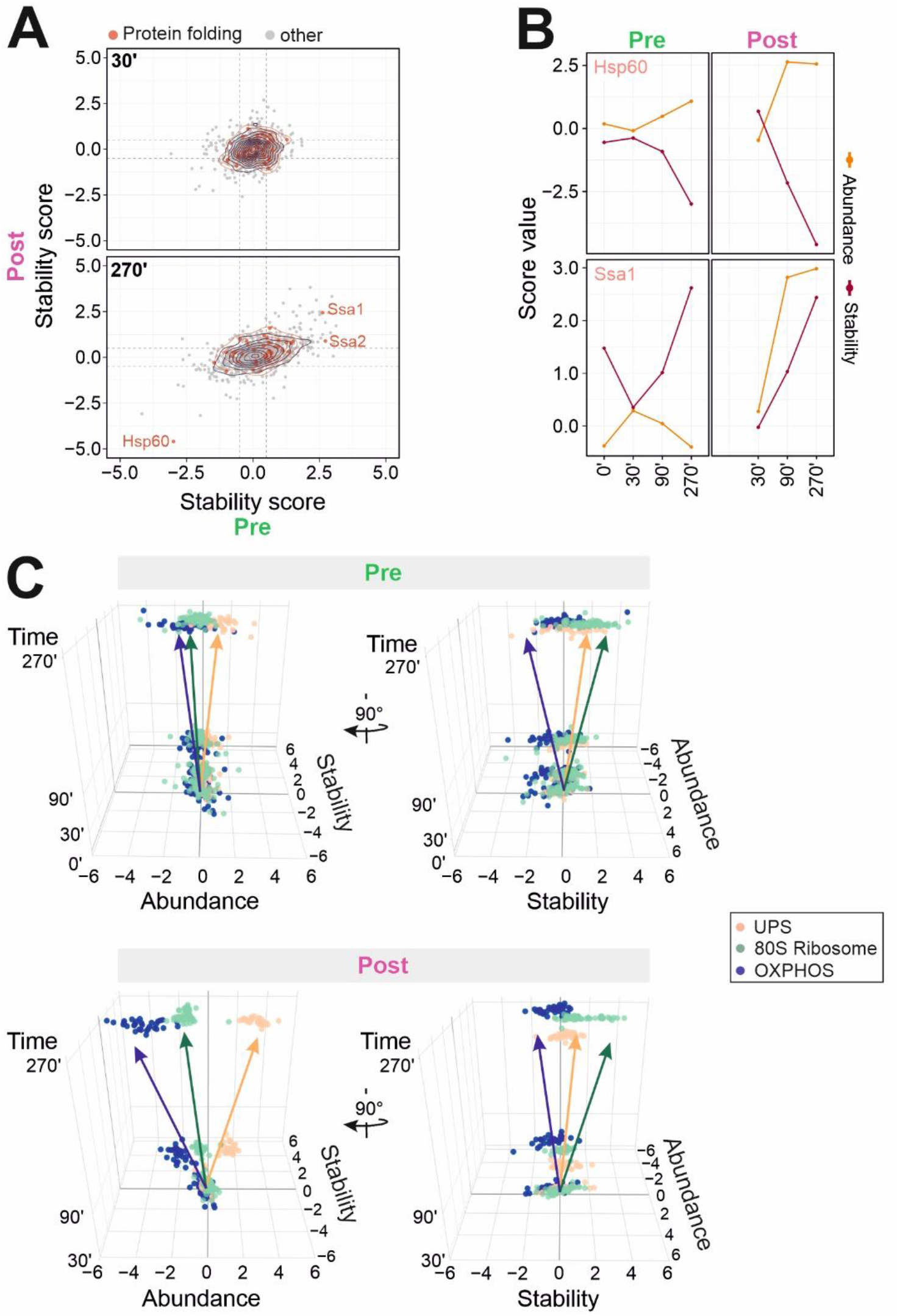
Cells react to mitoprotein-induced stress with complex adaptations of protein abundance and stability. **A**. Stability changes (clogger vs. control) of preexisting (x-axis) and newly synthesized proteins (y-axis) were correlated. Red circles show the isobaric distribution of chaperones. Black circles show the distribution of the whole proteome. **B**. The stress-induced abundance and stability changes are exemplarily shown for Hsp60 and Ssa1. **C**. Shown are the distinct functional groups of the UPS, the 80S ribosome and the OXPHOS system. For every protein of these groups, the stability and the abundance scores at the respective timepoint are shown in a 3D representation. Please note, that different groups of proteins follow individual trajectories in the stress response networks.

## Discussion

Cells adapt to different environmental conditions by remodeling of their proteome. These responses can provide resistance to extreme conditions as they arise from different types of stress situations. They are highly complex and affect many if not most proteins to different degrees. Still, on the first glance, cellular responses to stress conditions are seemingly similar (slower growth rates, induction of chaperones, increased proteasomal activity etc.) even though they need to resolve problems caused by protein-specific or locally defined perturbations. Transcriptomics and proteomics proved to serve as very powerful techniques to study genome-wide reactions in gene expression as well as in protein synthesis.

We have developed ppTPP, a strategy that combines pulsed SILAC and 2D-TPP with chemical labelling by tandem mass tags. Differentiating from previous methods, ppTPP is able to analyze the abundance and the thermal stability of mature and freshly synthesized proteins of multiple different treatment conditions in a single mass spectrometry-based experiment. Quantification of proteins already synthesized before onset of stress allows to detect changes in degradation, whereas *de novo* synthesized proteins allow to detect effects on posttranscriptional regulation. Furthermore, we are able to differentiate structural and hence functional changes between proteins synthesized before and after onset of stress. We discovered group-specific, as well as individual protein thermal stability variations which depend *inter alia* on the maturation and posttranslational modification of proteins and represent an additional layer of global proteome adaptation. In a nutshell, mitoprotein-induced stress generally leads on the level of abundance to (i) reduced protein synthesis including the repression of OXPHOS genes as well as the induction and increased assembly of the proteasome, (ii) increased protein degradation, and (iii) group-specific synergy between synthesis and degradation. On the level of thermal stability, our data could demonstrate that during stress conditions (i) newly synthesized proteins, prominently including mitochondrial proteins, show a higher vulnerability than non-mitochondrial proteins, (ii) many proteins of the cytosolic ribosome but not of the mitochondrial ribosome increased their stability, (iii) the chaperone network undergoes a remodeling to boost the cellular resistance against misfolded proteins, and (iv) diverse functional complexes, including histones and vesicle transport alter their stability but not their abundance to adapt to surrounding stress conditions. Our results with the higher vulnerability of nascent polypeptides can be explained by their compromised maturation. Since the melting temperature depends on the protein context (Mateus *et al*., 2020), the detected stability changes can indicate stress-induced alterations of posttranslational modifications, proteolytic cleavages, folding, complex formation or the localization of the protein. Whereas responses on the level of transcription often affect many genes in group-specific patterns (since these genes share regulatory elements such as the heat shock element HSE, the proteasome-associated control element PACE, the pleiotropic drug resistance element PDRE etc.), posttranscriptional regulation differentially affects individual proteins, for example by posttranslational modifications.

Our ppTPP analysis identified the complex pattern of stability changes in many individual proteins. These changes were independent of the changes in protein abundance and represent an additional level of regulation. The overall independence of the structural changes to the abundance highlights the need of the cell for a fast and effective reaction to adapt to stress conditions. This can be demonstrated with the mitochondrial protein Mge1. Clogger expression renders mature Mge1 unstable. Mge1 is the mitochondrial nucleotide exchange factor of the matrix Hsp70 protein and essential to convert Hsp70 into its ATP-bound form (Laloraya *et al*, 1994; Slutsky-Leiderman *et al*, 2007; Westermann *et al*, 1995). Both for the yeast Mge1 as well as for its human homologs GrpEL1/2 it was shown that oxidative stress conditions switch these proteins from an active soluble state into amyloid-type aggregates (Karri *et al*, 2019; Srivastava *et al*, 2017). It was proposed that this conversion serves as safety mechanism to switch off the mitochondrial import motor upon stress conditions (Craig, 2018; Michaelis *et al*, 2022; Mokranjac, 2020). Our proteome-wide analysis supports this structural and thereby functional change of Mge1, suggesting that clogger induction impairs ATP consumption by the import motor. Additionally, recent work demonstrated that the redistribution of mature mitochondrial Hsp70, which is involved in protein folding and import, is able to regulate the efficiency of protein import (Banerjee *et al*, 2022; Michaelis *et al*., 2022). With our ppTPP approach, we were able to detect such maturation-specific structural changes on a proteome-wide scale, which reveals new insight about protein functions and associations. Apparently, many different cellular complexes changed predominantly on the stability rather than on the abundance level when mitochondrial protein import was inhibited. Previous studies have shown that protein thermal stability often arises from changes in post-translational modifications or from altered binding partners of these proteins (Becher *et al*., 2018). The observed stability change of histones suggests that clogger induction directly or indirectly impacts on the chromatin structure in the nucleus. This is consistent with the recent observation that clogger expression arrests the cell cycle at the entry into S phase (Krämer *et al*., 2022).

Interestingly, we noticed an increase in stability of Hsp104 upon clogger expression, which is compatible with an increased substrate binding to this disaggregase. Hsp104 and the small heat shock protein Hsp42 control the formation of protein granules in the yeast cytosol which transiently store misfolded proteins during stress conditions (Böckler *et al*, 2017; Grousl *et al*, 2018; ka chaNowicka *et al*., 2021; Miller *et al*, 2015; Shakya *et al*., 2021; Xiao *et al*., 2021). Our observations suggest that cells form such Hsp104-bound granules also during mitoprotein-induced stress conditions, in consistence with recent observations (ka chaNowicka *et al*., 2021; Krämer *et al*., 2022; Xiao *et al*., 2021).

With the ppTPP strategy used in this study we have performed a multidimensional, time-resolved, proteome-wide abundance and stability profiling during proteotoxic stress conditions on basis of the mitoprotein-induced stress response. Our insights illustrate the challenge of the cell to orchestrate a network of qualitative and quantitative changes. We are confident that the ppTPP method described here will proof to be a powerful and widely used tool to study cellular responses to different types and amplitudes of stress conditions in the future.

## Methods

### Strains and growth conditions

All strains used in this study were derived from YPH499 (MATa *ura3 lys2 ade2 trp1 his3 leu2 arg4*)(Sikorski & Hieter, 1989) and were grown at 30°C in minimal synthetic respiratory medium (SLac) containing 0.67% (w/v) yeast nitrogen base and 2% lactate as carbon source. To induce the clogger from the *GAL1* promoter, cells were shifted to SLac medium containing 0.5% galactose.

### Growth Assays

Growth curves were performed in a 96 well plate, using the automated ELx808™ Absorbance Microplate Reader (BioTek®). The growth curves started in SLac medium without or with 0.5% galactose at OD_600_ 0.1 and the OD_600_ was measured every 10 min for 72 h at 30°C. The experiment was performed in triplicates and the mean was calculated and plotted in R. Standard deviation for every measurement is shown.

For drop dilution assays an OD_600_ of 1 was harvested from cultures grown under non-inducing conditions during exponential growth phase, washed with sterile water and a serial 1:10 dilution was done. From each dilution, 3 μl were dropped on the SLac plates containing (inducing) or lacking (non-inducing) 0.5% galactose followed by incubation at 30°C. The growth was documented after different days.

### Pulsed SILAC

Yeast culture of an arginine and lysine auxotroph yeast strain YPH499 *Δarg4* with clogger or cytosolic DHFR plasmid was grown in synthetic minimal medium containing 2% lactate as carbon source and ‘light’ isotopes of arginine (Arg0, ^12^C_6_/^14^N_4_) and lysine (Lys0, ^12^C_6_/^14^N_2_). In mid-logarithmic growth phase (OD 0.6 – 0.8), the medium was removed by centrifugation (5 min, 5,000g), cells were washed once in medium not containing arginine or lysine and resuspended in medium containing ‘heavy’ arginine (Arg10, ^13^C_6_/^15^N_4_) and lysine (Lys8, ^13^C_6_/^15^N_2_) plus 0.5% Gal to induce the clogger or the cytosolic DHFR control. Samples of 50 OD_600_*ml were collected by centrifugation (5 min, 5,000g) before induction (0 h) or after induction for 30, 90 and 270 min.

### Thermal proteome profiling and sample preparation

For the TPP experiment harvested cells from pulsed SILAC shift were used directly. Cells were washed with PBS, resuspended in the same buffer in volume equal to OD_600_ 0.1 and 20 μl were aliquoted to eleven wells of a PCR plate. After centrifugation (5 min, 4,000g), the plate was subjected to a temperature gradient (37°C, 37.8°C, 40.4°C, 44°C, 46.9°C, 49.8°C, 52.9°C, 55.5°C, 58.6°C, 62°C, 66.3°C) for 3 min in a PCR machine, followed by 3 min at room temperature. Cells were lysed with 30 μl lysis buffer [50 mg/ml zymolyase, 0.8% NP-40, 1× protease inhibitor, 1× phosphate inhibitor, 0.25 U/μl benzonase, and 1 mM MgCl_2_ in PBS] for 30 min shaking (200 rpm) at 30°C. Three freeze-thaw cycles (freezing in liquid nitrogen, followed by 1 min at 25°C in a PCR machine and vortexing). The plate was then centrifuged (5 min, 2,000g) to remove cell debris, and the supernatant was filtered at 500g for 5 min at 4°C through a 0.45-μm 96-well filter plate to remove protein aggregates. 4 μl of the flow through were subjected to a protein determination (Pierce BCA Protein Assay (Thermo Scientific, #23225). 25 μl of the remaining flow-through were mixed 1:1 with 2x sample buffer (180 mM Tris pH 6.8, 4% SDS, 20% glycerol, 0.1 g bromophenol blue). The 37°C samples were diluted with 1x sample buffer to achieve a protein concentration of 1 mg/ml. An equal volume of 1x sample buffer was added to all other samples of the same condition. Samples were stored at −80°C.

In-solution digests were performed as previously described (Hollmann *et al*, 2020): 10 μg of each lysate were subjected to an in-solution tryptic digest using a modified version of the Single-Pot Solid-Phase-enhanced Sample Preparation (SP3) protocol (Hughes *et al*, 2014; Moggridge *et al*, 2018). Lysates were added to Sera-Mag Beads (Thermo Scientific, #4515-2105-050250, 6515-2105-050250) in 10 μl 15% formic acid and 30 μl of ethanol. Binding of proteins was achieved by shaking for 15 min at room temperature. SDS was removed by 4 subsequent washes with 200 μl of 70% ethanol. Proteins were digested with 0.4 μg of sequencing grade modified trypsin (Promega, #V5111) in 40 μl Hepes/NaOH, pH 8.4 in the presence of 1.25 mM TCEP and 5 mM chloroacetamide (Sigma-Aldrich, #C0267) overnight at room temperature. Beads were separated, washed with 10 μl of an aqueous solution of 2% DMSO and the combined eluates were dried down.

Peptides were reconstituted in 10 μl of H_2_O and reacted with 80 μg of TMT10plex (Thermo Scientific, #90111) (Werner *et al*, 2014) label reagent dissolved in 4 μl of acetonitrile for 1 h at room temperature. Here, different conditions of one temperature point were combined in a single TMT-experiment (see Fig. 1A). Excess TMT reagent was quenched by the addition of 4 μl of an aqueous solution of 5% hydroxylamine (Sigma, 438227). Peptides were mixed, subjected to a reverse phase clean-up step (OASIS HLB 96-well μElution Plate, Waters #186001828BA) and subjected to an off-line fractionation under high pH condition (Hughes *et al*., 2014).

The resulting 12 fractions were then analyzed by LC-MS/MS on an Orbitrap Fusion Lumos mass spectrometer (Thermo Scientific) as previously described (Sridharan *et al*, 2019). To this end, peptides were separated using an Ultimate 3000 nano RSLC system (Dionex) equipped with a trapping cartridge (Precolumn C18 PepMap100, 5 mm, 300 μm i.d., 5 μm, 100 Å) and an analytical column (Acclaim PepMap 100. 75 × 50 cm C18, 3 mm, 100 Å) connected to a nanospray-Flex ion source. The peptides were loaded onto the trap column at 30 μl per min using solvent A (0.1% formic acid) and eluted using a gradient from 2 to 40% Solvent B (0.1% formic acid in acetonitrile) over 2h at 0.3 μl per min (all solvents were of LC-MS grade). The Orbitrap Fusion Lumos was operated in positive ion mode with a spray voltage of 2.4 kV and capillary temperature of 275 °C. Full scan MS spectra with a mass range of 375–1500 m/z were acquired in profile mode using a resolution of 120,000 (maximum fill time of 50 ms or a maximum of 4e5 ions (AGC) and a RF lens setting of 30%. Fragmentation was triggered for 3 s cycle time for peptide like features with charge states of 2–7 on the MS scan (data-dependent acquisition). Precursors were isolated using the quadrupole with a window of 0.7 m/z and fragmented with a normalized collision energy of 38. Fragment mass spectra were acquired in profile mode and a resolution of 30,000 in profile mode. Maximum fill time was set to 64 ms or an AGC target of 1e5 ions). The dynamic exclusion was set to 45 s.

Acquired data were analyzed using IsobarQuant (Franken *et al*., 2015) and Mascot V2.4 (Matrix Science) using a reverse UniProt FASTA Saccharomyces_cerrevisiae_database (UP000002311) including common contaminants. In order to distinguish between newly synthesized SILAC-labelled (heavy) and non-labelled (light) proteins, two separate Mascot searches were conducted. (1) The following modifications were taken into account for the identification of mature (light, pre induction) proteins: Carbamidomethyl (C, fixed), TMT10plex (K, fixed), Acetyl (N-term, variable), Oxidation (M, variable) and TMT10plex (N-term, variable). (2) For the analysis of peptides derived from newly synthesized isotope-labelled proteins (heavy), the following modifications were considered as previously described (Määttä *et al*, 2020): Carbamidomethyl (C, fixed), Label: 13C(6)15N(4) (R, fixed), TMT10plexSILAC (K, fixed; composition: 13C(10)15N(3)C(2)H(20)N(−1)O(2)), Acetyl (Protein N-term, variable), Oxidation (M, variable), TMT10plex (N-term, variable).

The mass error tolerance for full scan MS spectra was set to 10 ppm and for MS/MS spectra to 0.02 Da. A maximum of 2 missed cleavages were allowed. A minimum of 2 unique peptides with a peptide length of at least seven amino acids and a false discovery rate below 0.01 were required on the peptide and protein level (Savitski *et al*, 2015).

The raw output files of IsobarQuant (protein.txt – files) were processed using the R programming language (ISBN 3-900051-07-0). Only proteins that were quantified with at least two unique peptides were considered for the analysis. Moreover, only proteins which were identified in two out of three mass spec runs in one of the lowest temperatures (37 °C or 37.8°C), minimum five out of 11 temperatures and identified in at least 8 mass spec runs over all temperatures were kept for the analysis. Raw reporter ion signals (‘signal_sum’ columns) were first cleaned for batch effects using limma (Ritchie *et al*, 2015) and further normalized using vsn, variance stabilization normalization (Huber *et al*, 2002). Different normalization coefficients were estimated for each Temperature and heavy or light experiments. Abundance and stability scores were calculated as indicated in Mateus et al. (Mateus *et al*., 2020). A clogger / control ratio was calculated for each temperature and SILAC label separately. The abundance score was estimated by calculating an average ratio of the first two temperatures (37 °C and 37.8 °C) for each replicate and SILAC label. The remaining ratios were then divided by the respective abundance average and summed up to calculate the stability score. Abundance and stability score were transformed into a z-distribution using the ‘scale’ function of R. Both scores were scaled separately for each SILAC label. In order to estimate the significance, abundance and stability scores for each replicate have been tested for difference using limma (Ritchie *et al*., 2015). The number of identifications in the different temperatures for each replicate has been used as a weight. The t-values (output of limma) were analyzed with the ‘fdrtool’ function of the fdrtool package (Strimmer, 2008) in order to extract p-values and false discovery rates (fdr - q-values).

Experiments were performed in n=3 independent biological replicates. For correlation the Pearson method was used. R= Pearson coefficient; p value <0.05 considered as significant. For distributional testing two-sided Kolmogorov-Smirnoff test was used. P values <0.05 were considered as significant. The used tests are indicated in the figure legends. For reasons of clarity, the density plots showing the distribution of the stability score were cut in order to concentrate on the main distribution rather than on the outlier proteins. The programming language R (https://www.r-project.org) was used to analyze the data. All figures generated were assembled in CorelDraw X7.

### Gene ontology enrichment analysis

Gene enrichment analysis was performed using GOrilla (Eden *et al*, 2009) based on GO terms ‘Biological Process’ and ‘Cellular compartment’. All quantified proteins found in the respective SILAC channel were used as background. A custom list was used for target proteins. Analysis was performed using two unranked lists of genes (target and background lists). Benjamini-Hochberg-corrected pvalues were used. Only the top results with a false discovery rate [FDR] < 5% were considered and shown.

### Localization analysis

Localization of a protein was defined by GO-term or custom lists. The list of proteins linked to ‘mitochondria’ was acquired using data from a previous study (Morgenstern *et al*, 2017). The list of proteins assigned to protein folding proteins (GO:0006457), Ubiquitin-Proteasome-system (GO:0000502) and 80S ribosome (GO:0022626, or custom list filtered for ‘Rpl’ or ‘Rps’ proteins) were collected using genome wide annotation for yeast, primarily based on mapping using ORF identifiers from SGD. Bioconductor package ‘org.Sc.sgdSGD’ (Bioconductor). List of proteins assigned to OXPHOS system was gathered using the data from (Boos *et al*., 2019). List of mitoribosomal proteins was generated using data from (Desai *et al*, 2017).

### Calculation of synthesis and degradation rates

To calculate protein synthesis rates, the increase in heavy peptide intensity over time (30’, 90’ and 270’) at 37°C corrected for batch-effects was used for analysis. For protein degradation rates, the decrease in light peptide intensity over time (0’, 30’, 90’ and 270’) at 37°C corrected for batch-effects was used for analysis. A statistical approach was used to study the protein synthesis rate (KS)/ degradation rate (KD). In this mathematical model it is assumed that proteins are synthesized/degraded exponentially along the time course. The ΔKS/ΔKD value was calculated by determining the ratio between KS or KD of the clogger and the control strain.

### Transcriptional data

For the correlation analysis between transcription and translation, transcriptional data from a previous study were used (Boos *et al*., 2019). For the analysis, the data generated 270’ after induction were used.

### Presentation of ribosomal 3D models

Densities were generated based on the atomic coordinates of a *S. cerevisiae* 80S Ribosome (PDB-6Q8Y)(Tesina *et al*, 2019) by simulating densities at 7 Å resolution using the ‘molmap’ functionality in UCSF Chimera (Pettersen *et al*, 2004). Simulated densities were visualized using UCSF ChimeraX (Goddard *et al*, 2018). Figures were assembled using Inkscape.

## Data availability

The protein MS datasets produced in this study are available in the PRIDE database with the ProteomeXchange identifier PXD037741

## Acknowledgements

We thank Andrea Trinkaus, Vera Nehr and Sabine Knaus for technical assistance. We thank André Mateus for help with establishing the TPP protocol in yeast. We thank Timo Mühlhaus for discussions. This study was financially supported by the European Research Council (ERC 101052639 MitoCyto to JMH), the Deutsche Forschungsgemeinschaft (HE2803/10-1 and GRK2737-STRESSistance to JMH), the Landesforschungsinitiative Rheinland-Pfalz BioComp (to FB and JMH), the Damon Runyon Cancer Research Foundation (DRG-2461-22 to FB) and the Joachim Herz Stiftung (to FB and CG).

## Author contributions

CG and FB constructed the strains, performed the biochemical, proteomic and yeast genetics experiments and optimized the conditions for the experiments. CG, FB and PH carried out the ppTPP experiment. FS and CG analyzed the data. SF and SP generated ribosomal models. FB, MMS and JMH supervised the study. JMH and CG wrote the manuscript, with all authors critically reviewing it.

## Supplemental Information

**Fig. S1.**
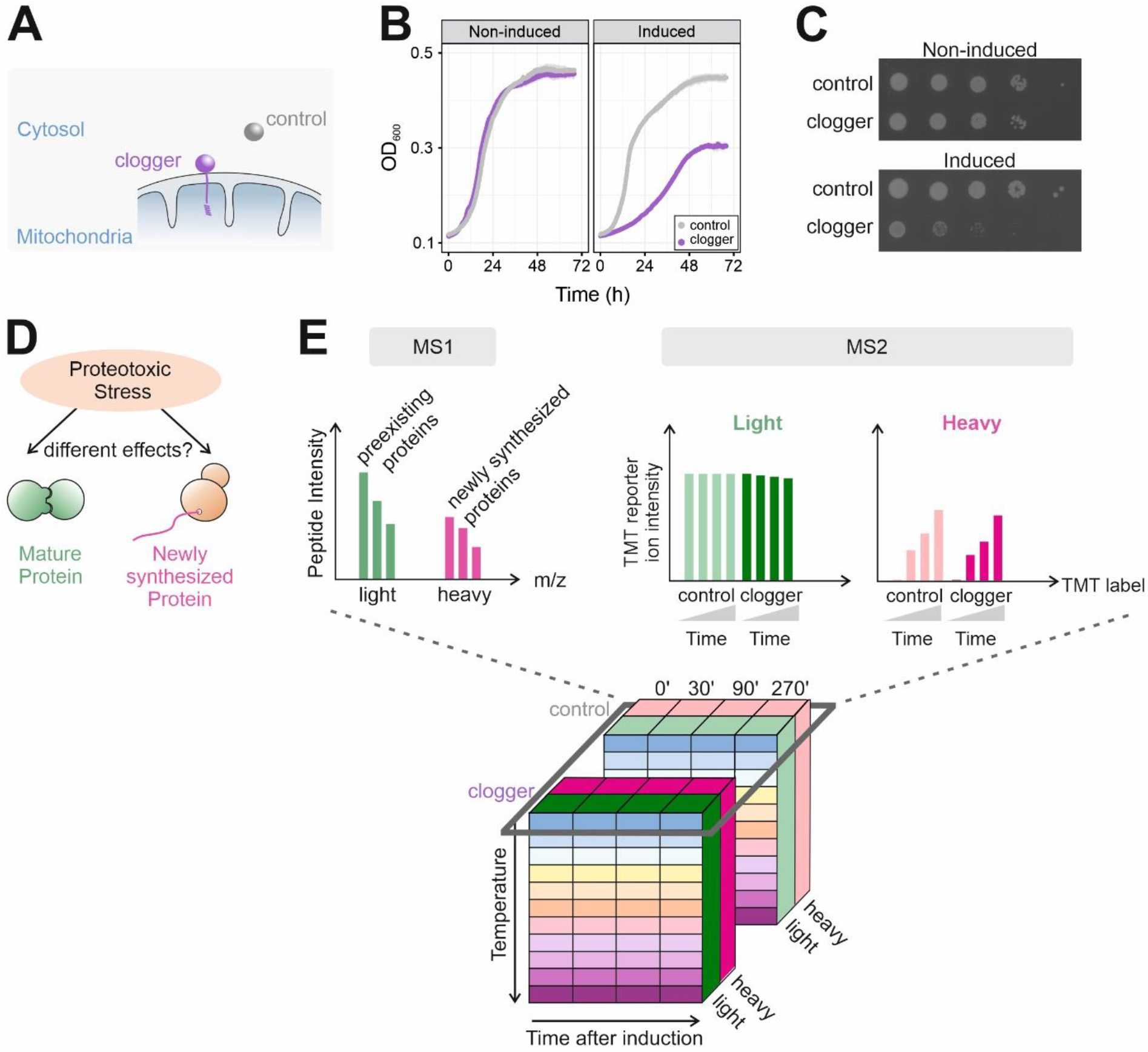
ppTPP as a multidimensional approach to investigate the effects of the mitochondrial-induced stress response. **A**. Schematic representation of the clogger and control protein. **B**. Clogger expression slows down cell growth. Cells were grown to mid-log phase in lactate medium. Then cells were used to inoculate cultures with lactate medium without (non-inducing) or with 0.5% galactose (inducing). The optical densities were continuously measured during growth at 30°C; n=3; error bars represent standard deviations. **C**. Cells were grown as in B and used to prepare ten-fold dilutions which were dropped on plates lacking (non-induced) or containing (induced) galactose. **D**. Pulsed SILAC allows the differentiation between preexisting and newly synthesized proteins. **E**. Cube representing the different dimensions of the ppTPP experiment. Dark grey box represents conditions that were measured together in one TMT experiment. Proteins in the soluble fractions were measured and quantified. Schematic representation of TMT reporter ions. MS1 allows to separate preexisting (light) from newly synthesized proteins (heavy). In MS2 peptides are identified and quantified based on TMT reporter ion intensities (TMT 8-plexing). Schematic representation of TMT reporter ions.

**Fig. S2.**
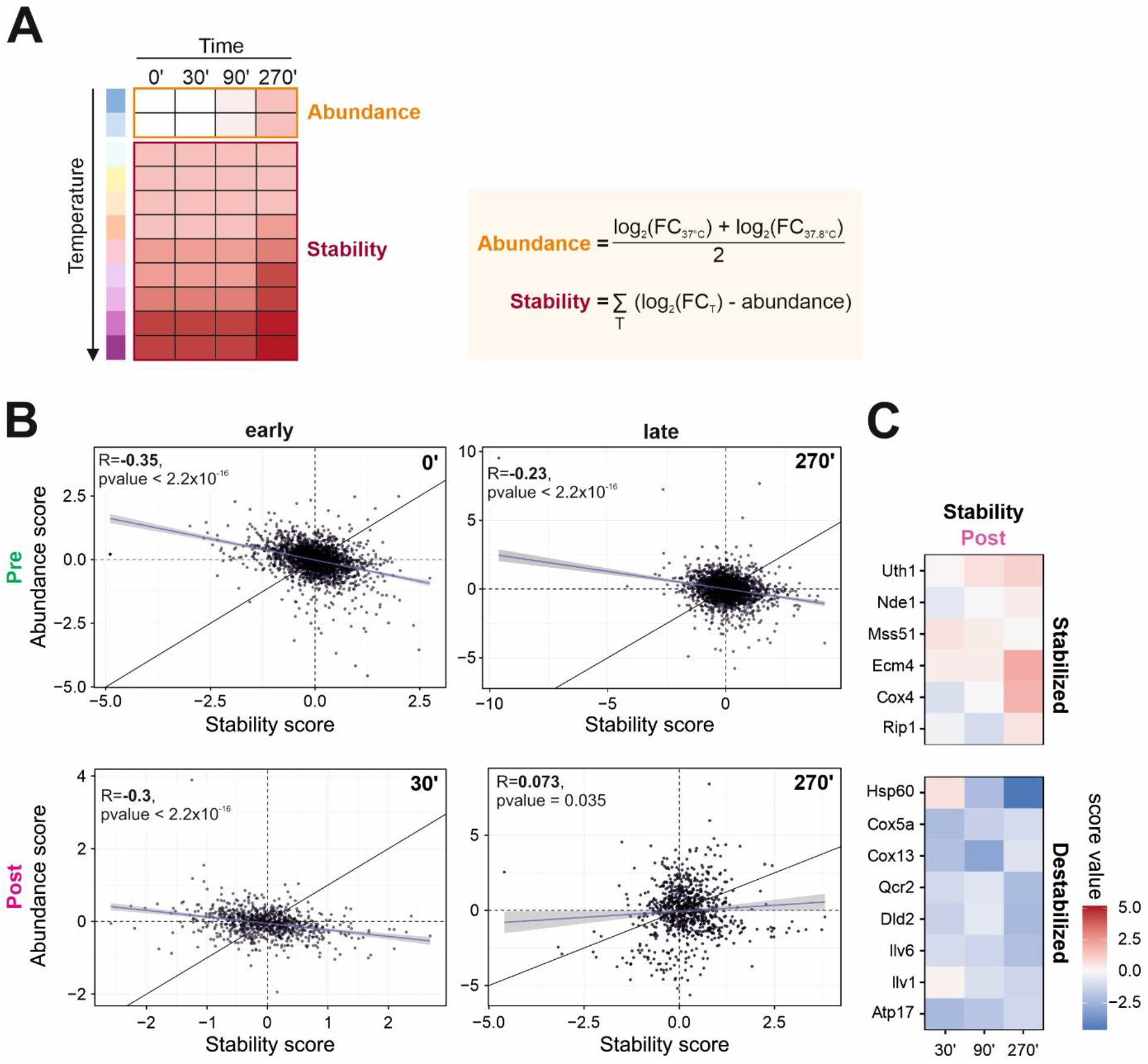
Stability of proteins clearly deviates from their abundance on a time-resolved basis. **A**. Calculation of abundance and stability score. The abundance score of each protein was calculated as the average log_2_ fold change of the two lowest temperatures. The thermal stability score of each protein was calculated by subtracting the abundance score from the log_2_-transformed fold changes of all temperatures and summing the resulting fold changes. **B**. Pearson correlation coefficients were calculated on basis of relative abundance and stability changes (clogger vs. control). Abundance poorly correlates with stability, especially for newly synthesized proteins after 270 min clogger induction. R, Pearson correlation coefficient. **C**. Zoomed inset of Fig. 2B demonstrates the advantage of ppTPP compared to using TPP exclusively. Shown are mitochondrial proteins, which stability is increased (upper panel) or decreased (lower panel) after induction of clogger.

**Fig. S3.**
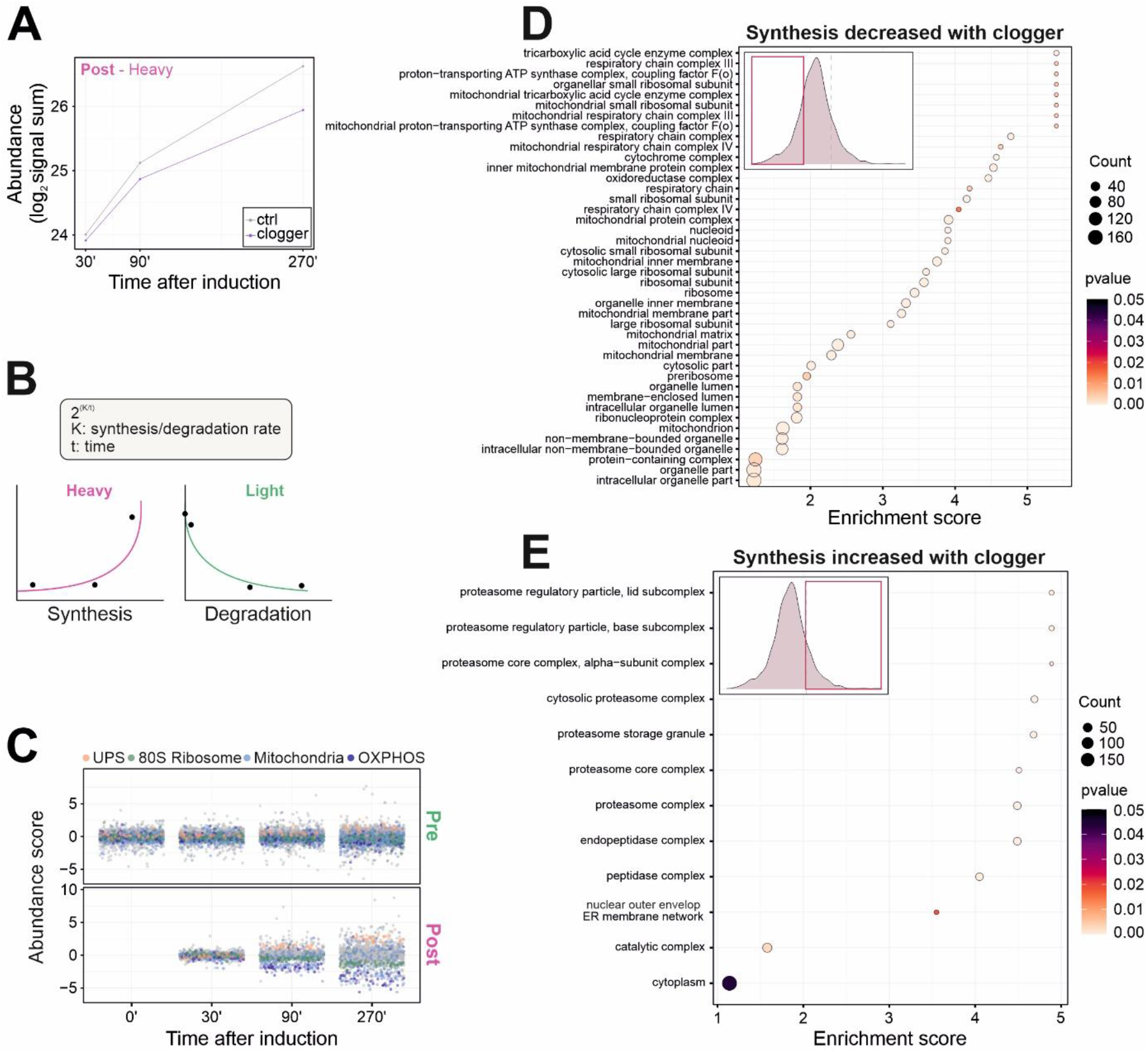
The clogger-induced stress response elicits a specific footprint on protein synthesis. **A**. Mean signal sums of newly synthesized (heavy) peptides in the absence and presence of clogger. **B**. Schematic representation of protein synthesis/degradation rate (K) calculation over time. For calculation of the synthesis rate (KS) signal intensities of heavy peptides were used, for degradation rate (KD) signal intensities of light peptides. **C**. Clogger induction elicits a characteristic footprint of the cellular proteome. Protein levels of freshly synthesized UPS proteins increase, 80S ribosomal proteins, mitochondrial proteins and the OXPHOS machinery are decreased over time. Shown is abundance score of clogger vs. control. **D**. GO enrichment analysis for newly synthesized proteins which accumulation upon clogger induction is prevented. Proteins of the lower 20% of all ΔKS values were used as target set and analyzed by using the GOrilla tool (http://cbl-gorilla.cs.technion.ac.il/) with all quantified proteins as background. The top results with a false discovery rate [FDR] < 5% are shown. **E**. GO enrichment analysis for newly synthesized proteins which synthesis upon clogger induction is increased. Proteins of the upper 20% of all ΔKS values were used as target set and analyzed by using the GOrilla tool (http://cbl-gorilla.cs.technion.ac.il/) with all quantified proteins as background. The top results with a false discovery rate [FDR] < 5% are shown.

**Fig. S4.**
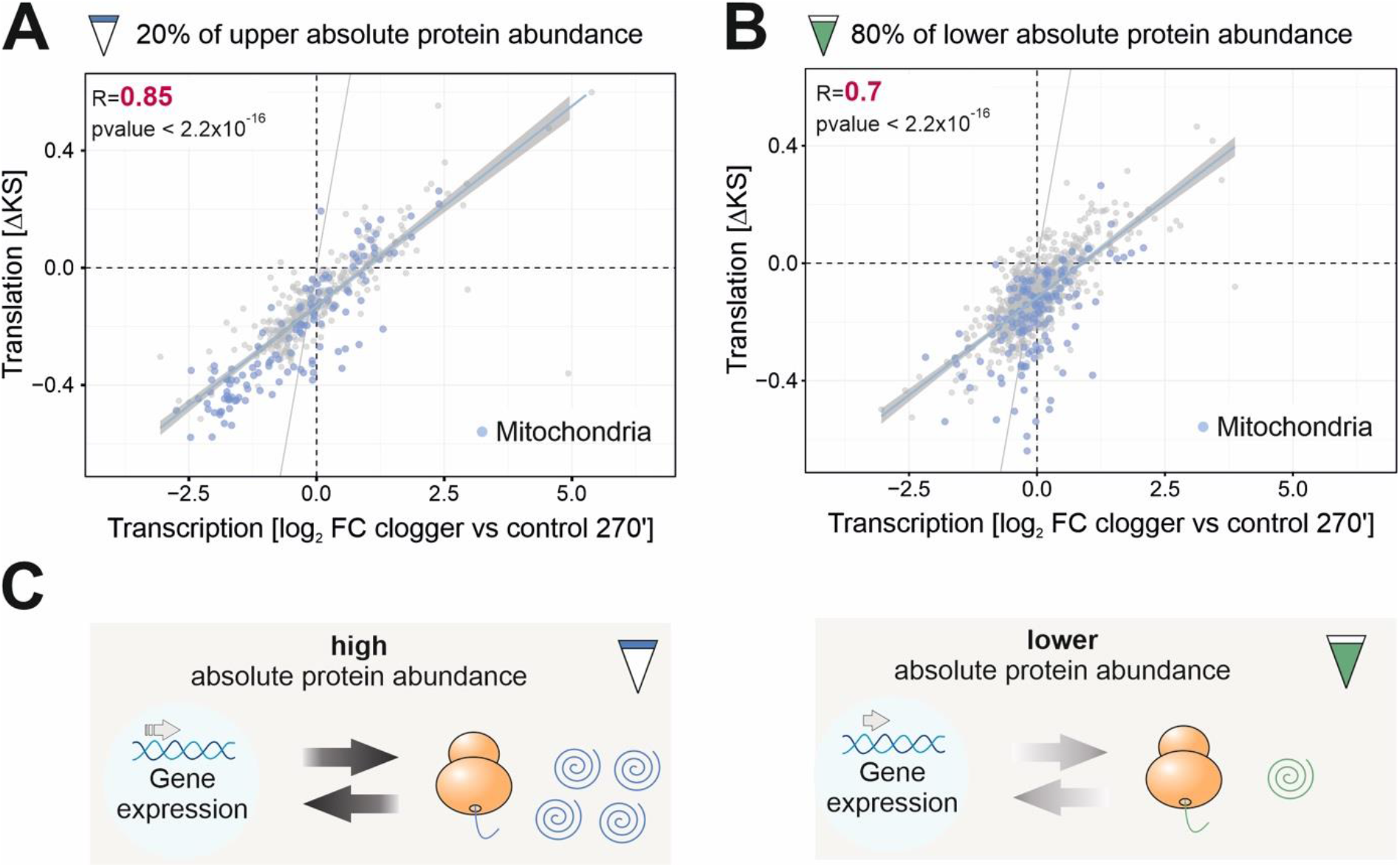
Changes in protein synthesis rate correlate with changes in mRNA abundance. **A**. Pearson correlation of transcription (Boos *et al*., 2019) with translation rate. Upper 20% of absolute protein abundance in the cell was used for analysis. Abundance values were obtained from previous quantifications (Morgenstern *et al*., 2017). Mitochondrial proteins are shown in blue. R, Pearson correlation coefficient. **B**. Pearson correlation of transcriptome data (Boos *et al*., 2019) with translation rate. Lower 80% of absolute protein abundance in the cell was used for analysis. Mitochondrial proteins are shown in blue. R, Pearson correlation coefficient. **C**. The correlation between transcription and translation is more pronounced for abundant proteins than for lower expressed proteins.

**Fig. S5.**
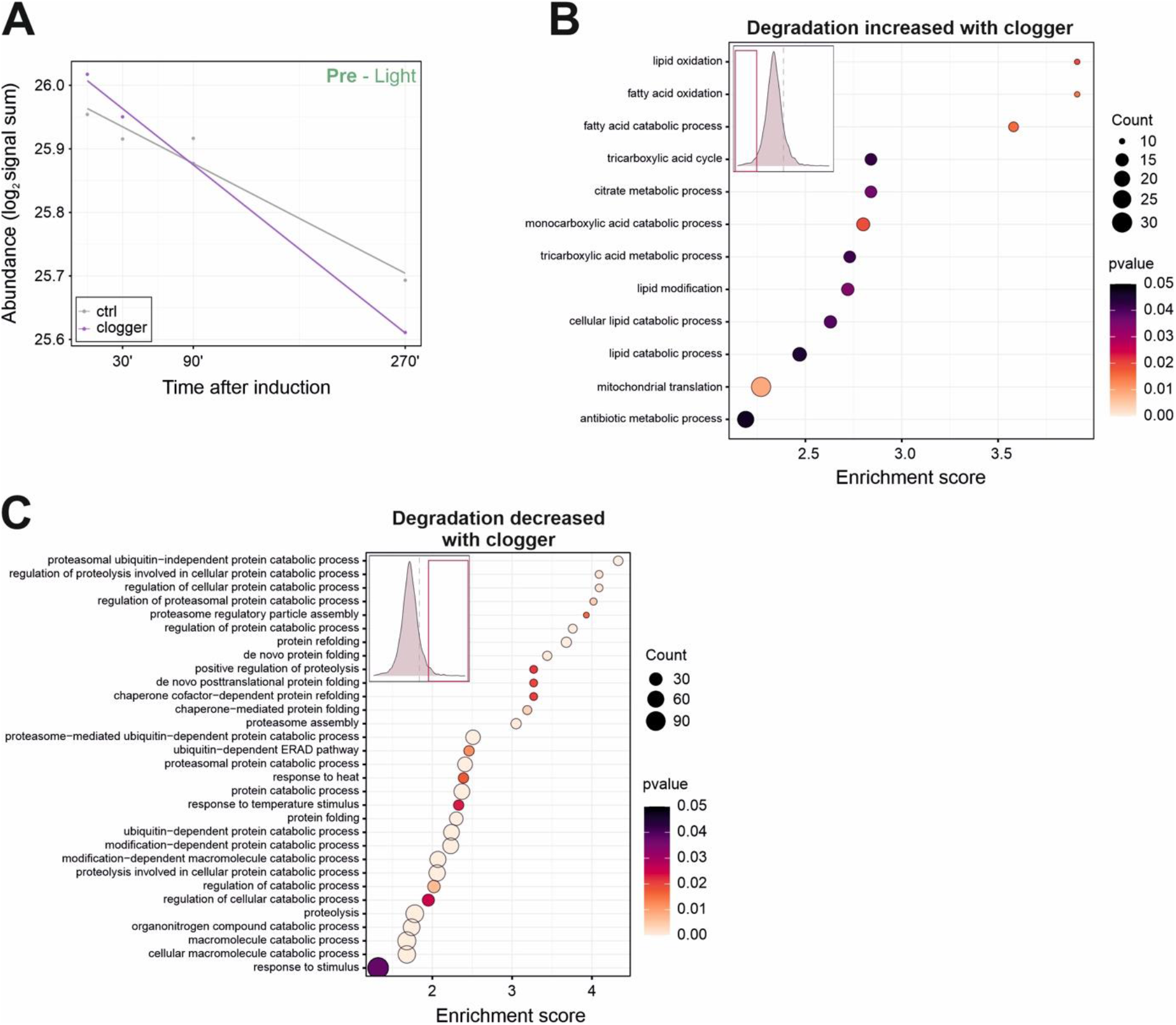
Clogger-induced stress leads to selective degradation of specific protein groups. **A**. Mean signal sums of preexisting (light) peptides in the absence and presence of clogger. **B**. GO-Enrichment analysis for preexisting proteins which degradation is induced upon clogger. The lower 20% of all ΔKD values were used as target set and analyzed by using the GOrilla tool (http://cbl-gorilla.cs.technion.ac.il/) with all quantified proteins as background. The top results with a false discovery rate [FDR] < 5% are shown. **C**. GO-Enrichment analysis for preexisting proteins which degradation is reduced upon clogger. The upper 20% of all ΔKD values were used as target set and analyzed by using the GOrilla tool (http://cbl-gorilla.cs.technion.ac.il/) with all quantified proteins as background. The top results with a false discovery rate [FDR] < 5% are shown.

**Fig. S6.**
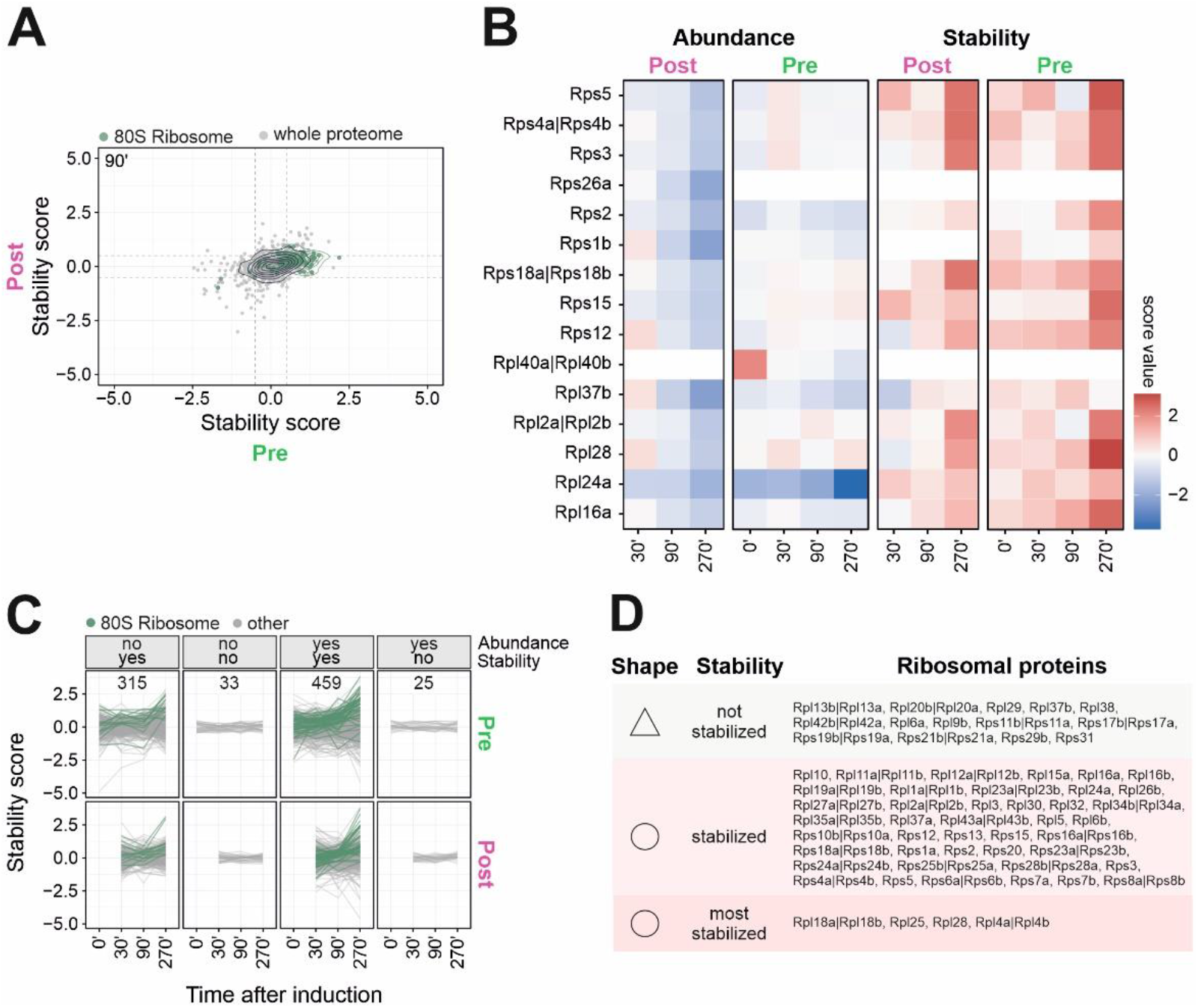
Many ribosomal proteins are strongly stabilized upon clogger induction. **A**. Clogger expression increases the thermal stability of many proteins both from preexisting as well as newly synthesized proteins of the 80S ribosome. Stability changes (clogger vs. control) of preexisting (x-axis) and newly synthesized proteins (y-axis) were correlated. Green circles show the isobaric distribution of 80S ribosomal proteins, which show an increased stability for both protein pools over time. Black circles show the distribution of the whole proteome. **B**. Heatmap showing the abundance and stability scores (clogger vs. control) of ribosomal proteins in the different samples. Heatmap from Fig. 2B was used and filtered for cytosolic ribosomal proteins. Data are demonstrating that upon clogger induction especially preexisting ribosomal proteins change their stability in comparison to their abundance. **C**. Classification for the different levels of protein regulation. Categories were generated on the basis of abundance and stability changes. ΔK value of the top or bottom 20% was defined as a change in abundance. A stability score (clogger vs. control) below −0.5 or above 0.5 was considered as change. Numbers indicate proteins per class. Cytosolic ribosomal proteins are shown in dark green many of which are represented in the group in which abundance does not considerably change, but stability does. See Table S3 for details. **D**. List which shows ribosomal proteins which increase their stability during clogger-induction. Stabilized is defined as stability score >0.5 for light- and heavy-encoded proteins, highly stabilized >2.9 for light-encoded proteins and > 0.5 for heavy-encoded proteins and no change as values below 0.5.

**Fig S7.**
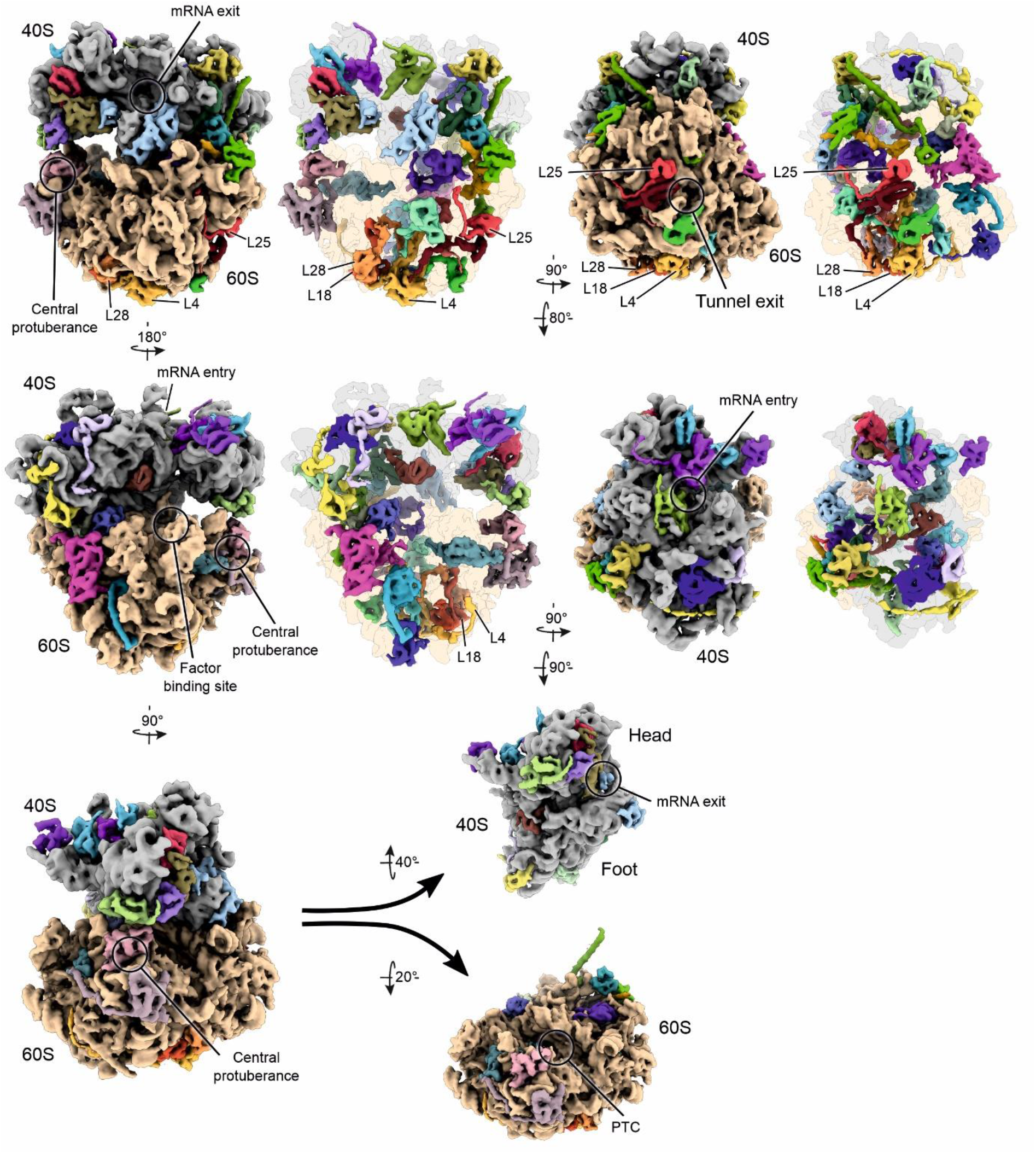
Clogger expression affects the thermal stability of many ribosomal proteins. Simulated 3D density of an *S. cerevisiae* 80S ribosome (based on PDB-6Q8Y)(Tesina *et al*., 2019) shown from various orientations. Moderately or most (L4, L18, L28, L25) stabilized ribosomal proteins colored individually.

**Fig. S8.**
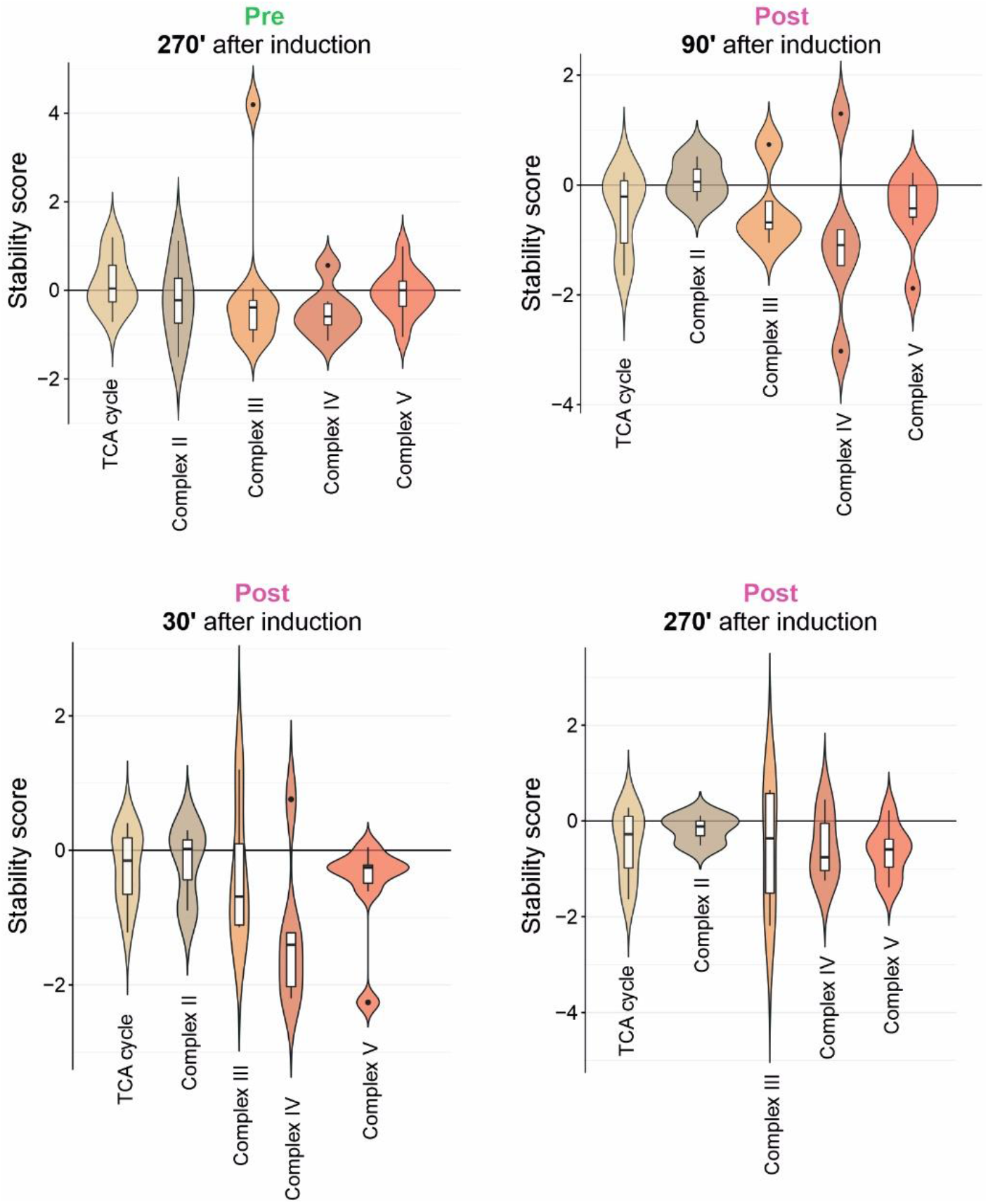
Stability of the subcomplexes of the OXPHOS is affected to varying degrees. Violin plots demonstrating the effect of the clogger on the different subcomplexes of the OXPHOS machinery for preexisting proteins and newly synthesized proteins. For the analysis, stability scores were calculated for the constituents of the functional groups indicated after different times of clogger expression.

**Fig. S9.**
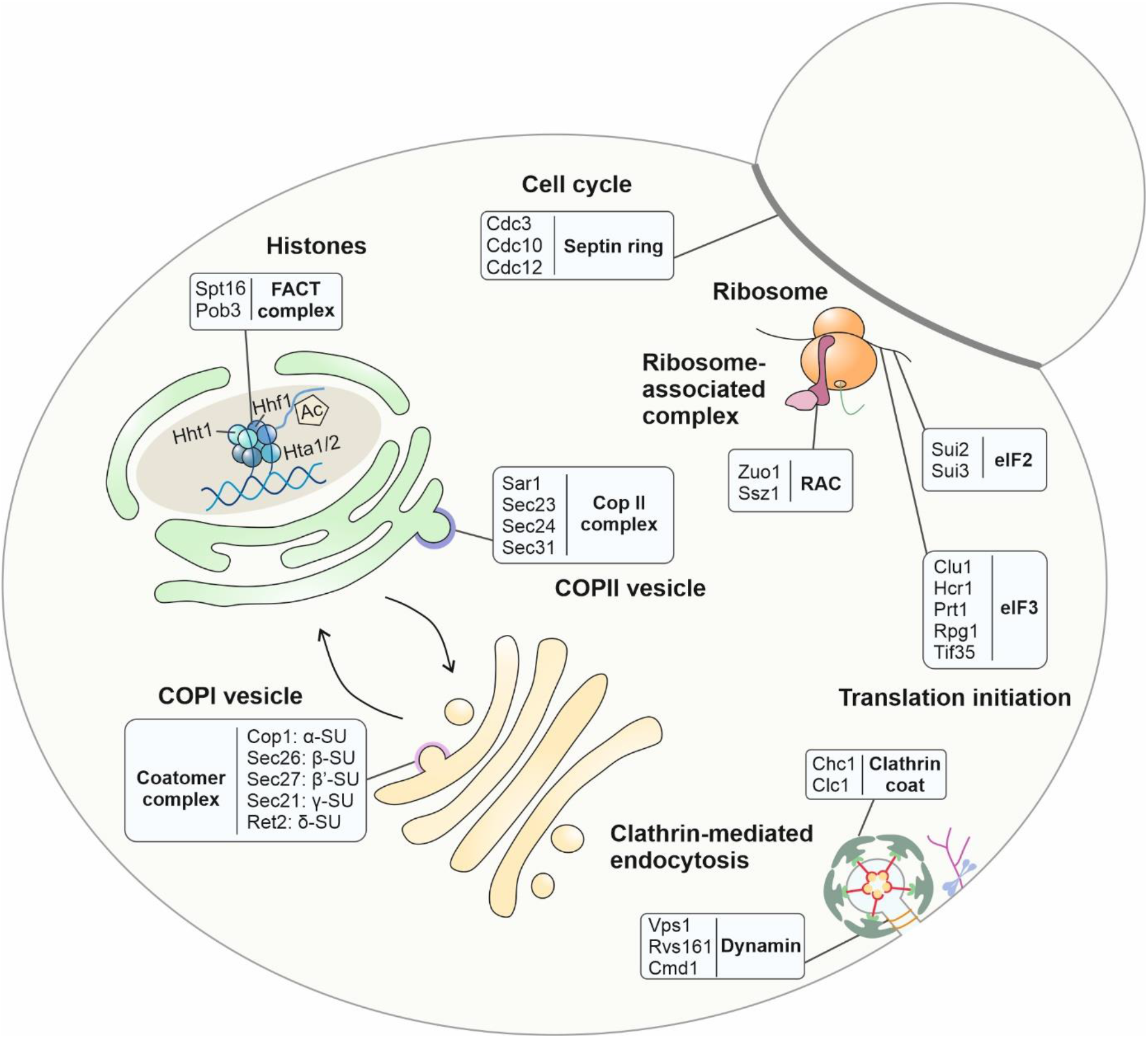
Clogger induction leaves a complex footprint on the stability of many cellular proteins without affecting their abundance. Examples of proteins that were not considerably changed in abundance (ΔK value within 20% and 80% range) but in stability (stability score below −0.5 or above 0.5) are shown. Proteins from complexes were taken into account, of which at least 60% are altered in stability but not in abundance. See Table S5 for details. Examples with clogger-provoked stability changes include: (i) for the histones H2A, H3 and H4 that form the nucleosome core complex for DNA packaging (Hta1/2, Hht1 and Hhf1) as well as the FACT (facilitates chromatin transcription) complex that remodels nucleosomes by histone acetylation (Ehara *et al*, 2022); (ii) many proteins of the cell cycle control system such as the components which form the septin ring (Cdc3, Cdc10, Ccd12); (iii) components of the ribosome-associated complex (RAC) as well as of the initiation factors of protein synthesis; as well as (iv) factors that control membrane trafficking including clathrin, dynamin as well as the coat proteins of COP I and COP II vesicles.

**Fig. S10.**
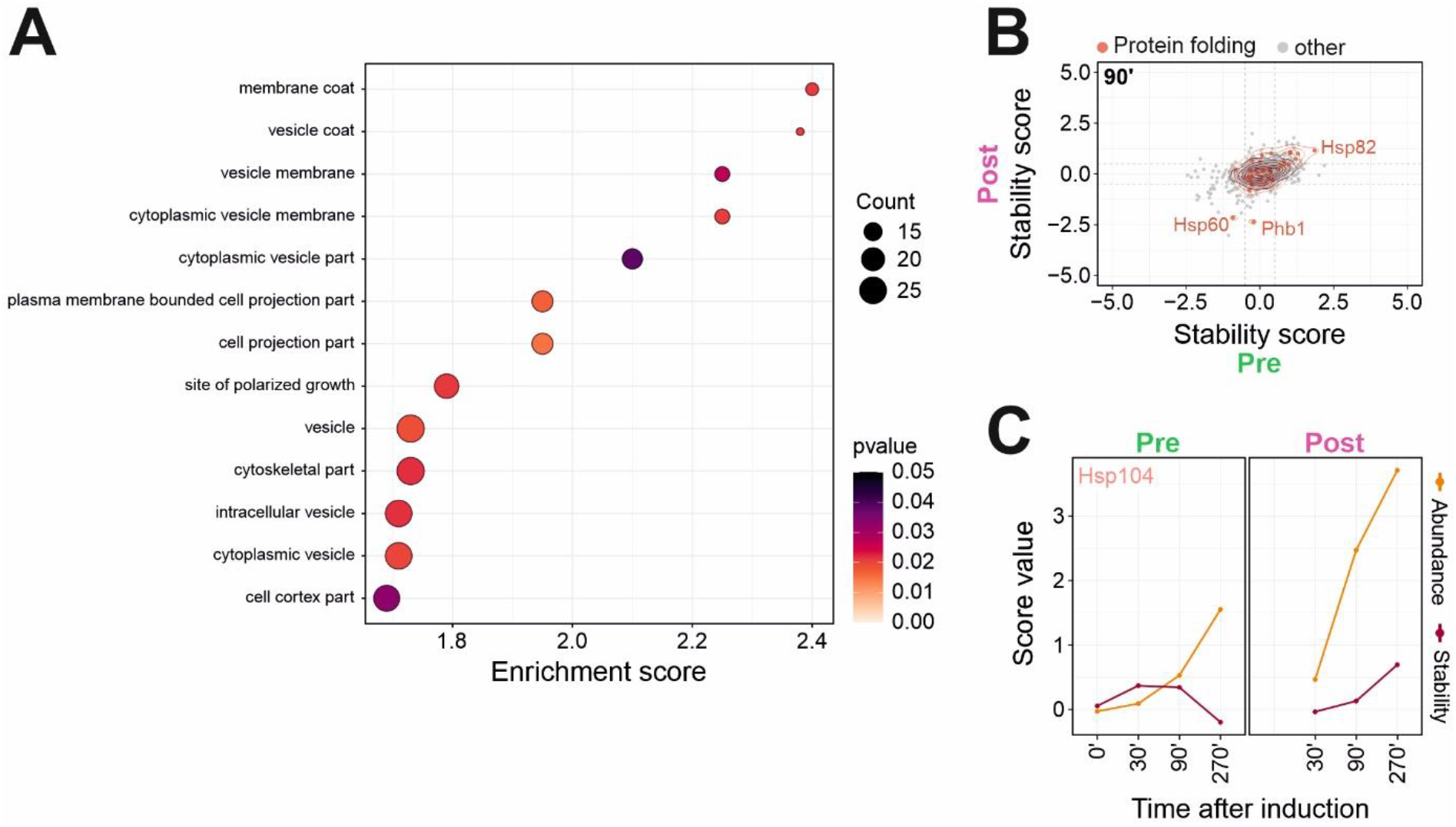
The stability of different chaperones reacts in a protein-specific manner on the inhibition of mitochondrial protein import. **A**. GO-Enrichment analysis for class of proteins in which abundance is not considerably changed but stability. All proteins of the class were used as target set and analyzed by using the GOrilla tool (http://cbl-gorilla.cs.technion.ac.il/) with all quantified proteins as background. The top results with a false discovery rate [FDR] < 5% are shown. **B**. The influence of the clogger on preexisting and newly synthesized protein folding proteins was analyzed. Red circles show the isobaric distribution of protein folding proteins, whereas black ones show the distribution of the entire proteome. **C**. The stress-induced abundance and stability changes are exemplarily shown for Hsp104.

## Notes

### Competing Interest Statement

The authors have declared no competing interest.

